# Compartmental Neuropeptide Release Measured Using a New Oxytocin Sensor

**DOI:** 10.1101/2022.02.10.480016

**Authors:** Tongrui Qian, Huan Wang, Peng Wang, Lan Geng, Long Mei, Takuya Osakada, Yan Tang, Alan Kania, Valery Grinevich, Ron Stoop, Dayu Lin, Minmin Luo, Yulong Li

**Affiliations:** State Key Laboratory of Membrane Biology, Peking University School of Life Sciences; Beijing, 100871, China; PKU-IDG/McGovern Institute for Brain Research; Beijing, 100871, China; National Institute of Biological Sciences (NIBS); Beijing, 102206, China; Chinese Institute for Brain Research; Beijing, 102206, China; Neuroscience Institute, Department of Psychiatry, New York University School of Medicine; New York, NY, USA; Center for Psychiatric Neurosciences, Department of Psychiatry, Lausanne University Hospital Center (CHUV); Lausanne, Switzerland; Department of Neuropeptide Research in Psychiatry, Central Institute of Mental Health, Medical Faculty Mannheim, University of Heidelberg; Mannheim, 68159, Germany; Tsinghua Institute of Multidisciplinary Biomedical Research (TIMBR), Tsinghua University; Beijing, 100084, China; Peking-Tsinghua Center for Life Sciences, Academy for Advanced Interdisciplinary Studies, Peking University; Beijing, 100871, China; National Biomedical Imaging Center, Peking University; Beijing, 100871, China

**Author notes:** These authors contributed equally to this work (Tongrui Qian, Huan Wang, Peng Wang).

## Abstract

As a peptide hormone and neuromodulator, oxytocin (OT) plays a critical role in a variety of physiological and pathophysiological processes in both the central nervous system and the periphery. However, the processes that regulate spatial OT release in the brain remain enigmatic. Here, we developed a genetically encoded GPCR activation-based (GRAB) OT sensor called GRAB_OT1.0_. Using this sensor, we directly visualized stimulation-induced OT release from specific compartments of OT neurons in acute brain slices, and discovered that N-type calcium channels predominantly mediate axonal OT release, while L-type calcium channels mediate somatodendritic OT release. In addition, we found that components in the fusion machinery of OT release differ between axon terminals versus somata and dendrites. Finally, we demonstrated the sensor responses to the activation of OT neurons in various brain regions *in vivo* and revealed region specific OT release during male courtship behavior. Taken together, these results provide key insights regarding the role of compartmental OT release in the control of physiological and behavioral functions.

## Main

Neuronal communication is typically described as a unidirectional process in which dendrites receive and integrate input information, and the soma transforms the signals into action potentials, which then propagate to the axon terminal to release neurotransmitters. However, in addition to classical axonal release, some neurochemicals, such as GABA^1^, dopamine^2^, and neuropeptides^3–5^, also undergo somatodendritic release to reciprocally modulate surrounding neurons and regulate important physiological functions^6, 7^. Evolutionarily ancient magnocellular neurosecretory cells (MNCs) in the paraventricular (PVN) and supraoptic (SON) nuclei of the hypothalamus, contributed extensively to the understanding of neurosecretory mechanisms, having been shown to release oxytocin (OT) or arginine-vasopressin (AVP) from both the axonal and somatodendritic compartments^5, 8, 9^.

OT is known to regulate a range of physiological processes in both the periphery and the central nervous system. In the mammalian brain, OT is produced by neurons located primarily in the PVN, SON, and accessory nuclei of the hypothalamus^10^. The OT synthesized in these brain regions is released into the blood circulation from the posterior pituitary to serve as a hormone, regulating parturition and lactation via oxytocin receptors (OTRs) that are robustly expressed in the uterus and mammary gland, respectively^11, 12^. Apart from projecting to the pituitary, OT neurons send axons throughout the brain where axon-released OT modulates food intake, fear, aggression, social, sexual, and maternal behaviors in rodents^13^, while somatodendritic OT release has been associated with autocrine functions during milk ejection and uterine contraction^14^. Coincident with its various physiological roles, altered regulation of OT signaling has been associated with various negative emotional states and conditions such as stress, social amnesia, autism spectrum disorder, and schizophrenia^15–18^.

Not only does compartmental OT release display distinct functions, somatodendritic OT release can also be primed by mobilization of intracellular Ca^2+^ which has no reported effect on axonal OT release^5^. Thus, OT is likely to be released independently from each compartment. However, the molecular mechanism underlying the compartmental control of OT release are largely unknown. The lack of sensitive, specific, and non-invasive tools to monitor OT dynamics at high temporal and spatial resolution imposes limitations on studying the mechanisms and functions of compartmental OT release. Existing methods used to measure OT release have inherent limitations. For example, the temporal and spatial resolution of microdialysis used to monitor extracellular OT in freely moving rats^19–21^ is too low to distinguish between axonal and somatodendritic release. Cell-based OT detection^22^ uses exogenous ‘sniffer cells’ expressing OTR to ‘sniff’ OT and reports it with increased intracellular Ca^2+^ level. This method, however, is limited by *ex vivo* setup as well as invasive and lacks stable spatial resolution due to the random distribution of ‘sniffer cells’. Finally, measuring downstream signals using the recently developed genetically encoded OT sensor OTR-iTango2 requires the co-expression of at least three components, as well as long-term light-induced activation of reporter gene expression^23^, making this approach relatively complicated and temporally underresolved.

To overcome these limitations, we developed a highly sensitive, OT-specific G protein‒coupled receptor (GPCR) activation‒based (GRAB) sensor. We used this sensor *ex vivo* to identify the mechanisms of OT release in distinct cellular compartments. We also imaged the sensor in behaving mice, and revealed region specific OT release during discrete aspects of mating behaviors in male mice. Together, this work expands the toolbox of genetically encoded sensors for neurotransmitters and neuromodulators^24–34^ to elucidate novel mechanisms of peptidergic signaling in the brain.

### Development and *in vitro* characterization of a GRAB_OT_ sensor

To measure the dynamics of extracellular OT with high temporal and spatial resolution, we designed a genetically encoded GRAB sensor specific for OT (GRAB_OT_). In this sensor, the circularly permutated GFP (cpGFP), which is flanked by linker peptides derived from the well-characterized GRAB_NE1m_ norepinephrine sensor^27^, serves as the fluorescent module, while the OT receptor functions as the ligand-recognition module (**Fig. 1a**). First, we systematically screened OTRs derived from ten vertebrate species (human, mouse, rat, bovine, sheep, pig, cat, chicken, monkey, and medaka) by inserting the fluorescent module into the third intracellular loop (ICL_3_) of each OTR with various insertion sites; we then measured each sensor’s change in fluorescence (ΔF/F_0_) in response to OT (**Fig. 1b**). We selected the bovine OTR‒based sensor (which we called GRAB_OT0.5_) for further optimization due to its large change in fluorescence intensity upon OT application. We then screened approximately 300 variants of GRAB_OT0.5_ with different ICL_3_ lengths, resulting in the optimized GRAB_OT1.0_ sensor (hereafter referred to as OT1.0) with the highest response to OT and specific membrane targeting (**Fig. 1c**). When we expressed OT1.0 in HEK293T cells, we found that bath application of OT induced a robust, stable response that was abolished by pretreating cells with the OTR antagonist Atosiban, but was not affected by the cholecystokinin B receptor antagonist YM022 (**Fig. 1d**). For use as a negative control, we also developed an OT-insensitive variant called OTmut, which traffics to the plasma membrane but does not respond to OT (**Extended Data Fig. 1a-c**).

**Fig. 1:**
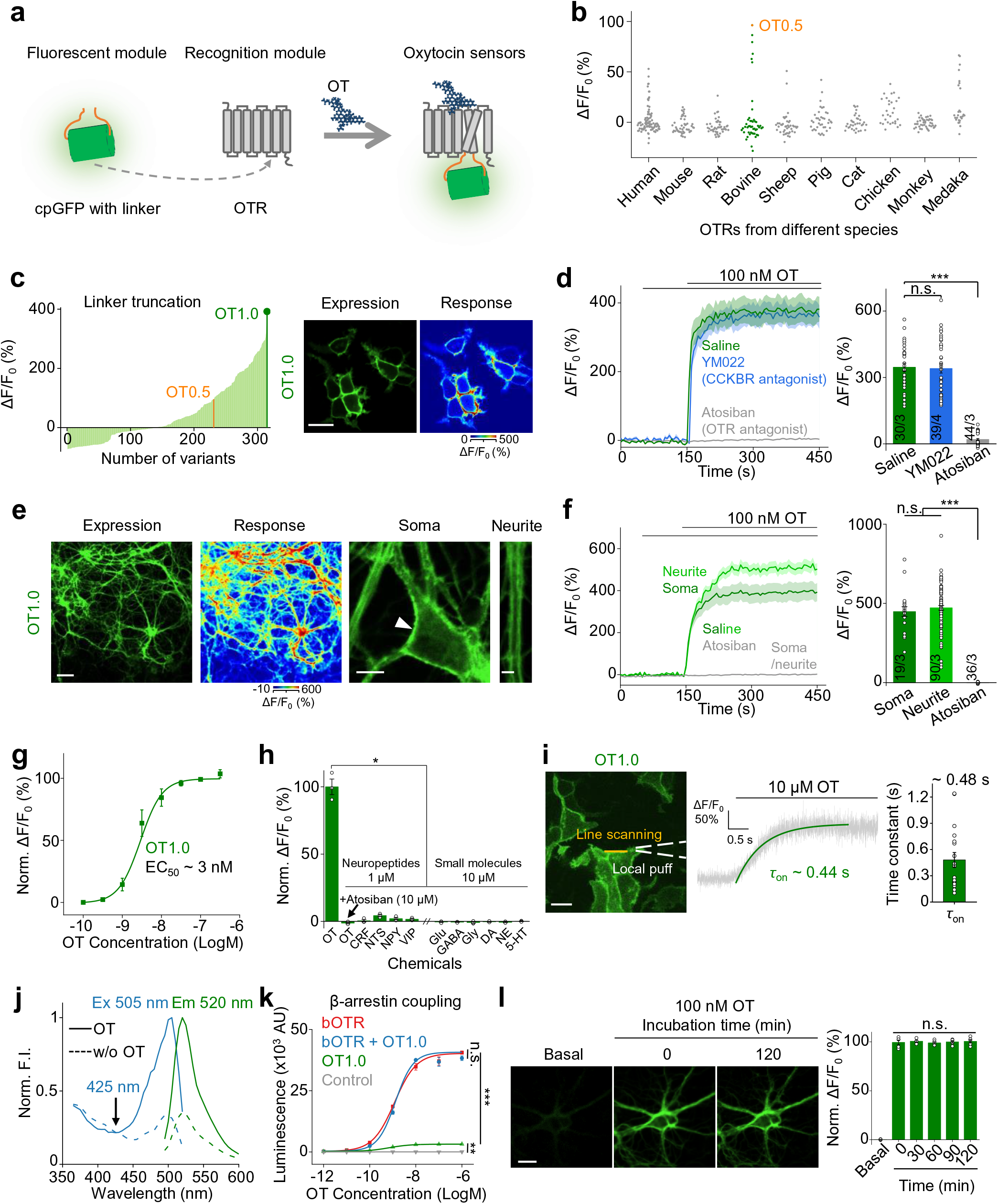
Development, optimization, and *in vitro* characterization of GRAB_OT_ sensors. **a.** Schematic diagram depicting the principle behind the GRAB_OT_ sensor, which contains cpGFP with a linker as the fluorescent module and the oxytocin receptor (OTR) as the recognition module. The third intracellular loop in the OTR is replaced with cpGFP connected with the linker, and OT binding activates the sensor, inducing an increase in fluorescence. **b.** Selection of a candidate sensor for further development by screening a wide range of OTRs cloned from the indicated species and expressed in HEK293T cells. The peak change in fluorescence relative to baseline (ΔF/F_0_) in response to 100 nM OT is shown, and each symbol represents a different sensor candidate in which cpGFP was inserted at a different site. The orange symbol indicates the OT0.5 candidate, which had the strongest response among all candidates. **c.** The candidate GRAB_OT0.5_ sensor was optimized by adjusting the length of the linker (left). The GRAB_OT1.0_ sensor used in this study was identified as having the highest response (ΔF/F_0_) among all variants tested. Also shown are representative images of OT1.0 expression and the response to 100 nM OT (right) in HEK293T cells. Scale bar, 20 μm. **d.** Example fluorescence traces (left) and summary data (right) of OT1.0-expressing HEK293T cells pre-incubated with saline (green), the CCKBR antagonist YM022 (10 μM, blue), or the OTR antagonist Atosiban (10 μM, gray); where indicated, 100 nM OT was applied. Saline: n=30 cells from 3 coverslips [30/3], YM022: n=39/4, Atosiban: n=44/3. **e.** Representative images of OT1.0 expression and the response to 100 nM OT (left) in cultured rat cortical neurons expressing OT1.0. Also shown are representative images of OT1.0 expression in the soma (arrowhead) and in a neurite. Scale bars represent 50 μm, 10 μm, and 5 μm for the expression/response, soma, and neurite images, respectively. **f.** Example fluorescence traces (left) and summary data (right) of OT1.0-expressing neurons pretreated with saline (neurite in green and soma in dark green) or 10 μM Atosiban (gray); where indicated, 100 nM OT was applied. Soma: n=19 ROIs from 3 cultures [19/3], neurite: n=90/3, Atosiban: n=36/3. **g.** Normalized dose–response curve of OT1.0-expressing neurons in response to the indicated concentrations of OT; n=4 cultures each with >30 cells. **h.** Summary of ΔF/F_0_ measured in OT1.0-expressing neurons in response to the indicated compounds applied at the indicated concentrations and normalized to the peak response measured in OT. CRF, corticotropin-releasing factor; NTS, neurotensin; NPY, neuropeptide Y; VIP, vasoactive intestinal peptide; Glu, glutamate; GABA, γ-aminobutyric acid; Gly, glycine; DA, dopamine; NE, norepinephrine; 5-HT, 5-hydroxytryptamine (serotonin). n=3 wells per group, with each well containing 200-400 neurons. **i.** Summary of the kinetics of the OT1.0 response. OT was locally puffed onto an HEK293T cell expressing OT1.0, and high-speed line scanning was used to measure the fluorescence response (left). The representative trace (middle) shows the change in OT1.0 fluorescence in response to 10 μM OT, and τ_on_ is summarized at the right; n=18 cells from 3 cultures. Scale bar, 25 μm. **j.** Excitation (Ex) and emission (Em) spectra of the OT1.0 sensor in the presence (solid lines) and absence (dashed lines) of 100 nM OT. **k.** β-arrestin coupling was measured using the Tango assay in HTLA cells expressing the bovine OTR (bOTR) alone, OT1.0 alone, bOTR and OT1.0, or no receptor (control) in the presence of the indicated concentrations of OT. n=3 wells each. **l.** Representative images (left) and summary (right) of the fluorescence change measured in OT1.0-expressing neurons in response to a 2-hour continuous application of 100 nM OT. n=5 cultures with >30 cells each. Scale bar, 20 μm. \**p*<0.05, \*\**p*<0.01, \*\*\**p*<0.001, and n.s., not significant.

When expressed in cultured rat cortical neurons, OT1.0 was present throughout the plasma membrane, including the soma and neurites (**Fig. 1e**). Upon application of a saturating concentration of OT, both the soma and neurites displayed a robust increase in fluorescence, which was prevented by pretreatment with Atosiban (**Fig. 1e, f**). We also measured the dose-response curve for OT1.0 expressed in neurons, with 3 nM OT eliciting the half-maximal effect (EC_50_ ≈ 3 nM; **Fig. 1g**). OT1.0 had high specificity for OT, whose EC_50_ value was ∼12 fold lower than that of AVP (**Extended Data Fig. 1c**), and had virtually no response to a wide range of other neurotransmitters and mammalian neuropeptides when expressed in cultured HEK293T cells (**Extended Data Fig. 1e**) or cortical neurons (**Fig. 1h**). Moreover, OT1.0 could be potentially used to monitor OT orthologs of other species, including vasotocin (a non-mammalian AVP-like hormone) and isotocin (OT homologue in fish), which had slightly higher EC_50_ than oxytocin (**Extended Data Fig. 1d**). Next, we examined whether the OT1.0 sensor has a response sufficiently rapid to capture OT applied using a local puffing system combined with high-speed line scanning. We found that OT1.0 has rapid response kinetics, with an average rise time constant (τ_on_) of 480 ± 84 ms (**Fig. 1i**). We then characterized the spectral properties of OT1.0 expressed in HEK293T cells before and after OT application, and measured the peak excitation (Ex) and emission (Em) wavelengths to be 505 nm and 520 nm, respectively (**Fig. 1j**).

To confirm that the OT1.0 sensor does not couple to the downstream signaling pathways (and therefore does not affect cellular physiology), we used the Tango GPCR assay^35^ to measure β-arrestin activation (**Fig. 1k**). When expressed in HTLA cells (an HEK293T-derived cell line expressing a tTA-dependent luciferase reporter and a β-arrestin2-TEV fusion gene), OT1.0 had a minimal β-arrestin coupling in response to OT; in contrast, the wild-type bovine OTR (bOTR) displayed a robust coupling (**Fig. 1k**). Importantly, OT1.0 does not interfere with signaling via the wild-type bOTR, as cells co-expressing OT1.0 and bOTR had the same response to OT as cells expressing OTR alone (**Fig. 1k**). Moreover, there is no detectable coupling from OT1.0 to the Gq-dependent calcium signaling in defined OT concentrations^11, 36^ compared to the wild-type bOTR (**Extended Data Fig. 2**). Finally, we found that the OT response measured in OT1.0-expressing neurons was extremely stable for up to 120 min, indicating that the sensor undergoes negligible internalization or desensitization and can be used for long-term imaging of OT (**Fig. 1l**). Taken together, these results confirm that the OT1.0 sensor can be used *in vitro* to measure OT release with high sensitivity, high specificity, and rapid kinetics.

### Axonal and somatodendritic OT release measured in acute brain slices

Having shown that our OT1.0 sensor can be used in cultured cells, we next examined whether it can be applied to monitor the endogenous OT release from oxytocinergic neurons in acute brain slices. We therefore injected adeno-associated viruses (AAVs) expressing OT1.0 (hSyn-OT1.0) and Cre-dependent excitatory Gq-DREADDs (EF1α-DIO-hM3Dq-mCherry) into the PVN of OT-Cre mice (**Extended Data Fig. 3a**). After 3 weeks (to allow sufficient sensor expression), we found that activation of oxytocinergic neurons by bath application of the hM3Dq ligand deschloroclozapine (DCZ)^37^ elicited a robust increase in OT1.0 fluorescence in the PVN, with a stronger response elicited by direct application of OT (**Extended Data Fig. 3b, c**). In contrast, DCZ elicited virtually no OT1.0 response in slices not expressing hM3Dq, while OT application still produced a peak response comparable to slices expressing hM3Dq (**Extended Data Fig. 3b-d**). Thus, OT1.0 can report the OT released from activated OT neurons.

We next tested whether OT1.0 can detect the dynamics of compartmental OT release in acute brain slices. PVN originating OT input to the ventral tegmental area (VTA) has been shown to regulate social behavior via OT/OTR-mediated modulation of dopaminergic neurons^38–40^. Therefore, we injected a virus expressing either OT1.0 or OTmut under the control of hSyn promotor into the VTA to visualize axonal OT release. After at least 3 weeks, acute sagittal brain slices containing the VTA region were prepared and used for two-photon imaging (**Fig. 2a**). We found that electrical stimulation delivered at 20 Hz evoked a progressively larger fluorescence increase in OT1.0, which was eliminated by treating slices with the OTR antagonist L368. Moreover, no response was measured in slices expressing OTmut (**Fig. 2b, c**). We also measured the kinetics of axonal OT release in response to 20, 50 and 100 pulses delivered at 20 Hz and estimated rise time (τ_on_) and decay time constants (τ_off_) of 1.3-3.2 s and 6.6-9.9 s, respectively (**Fig. 2d**).

**Fig. 2:**
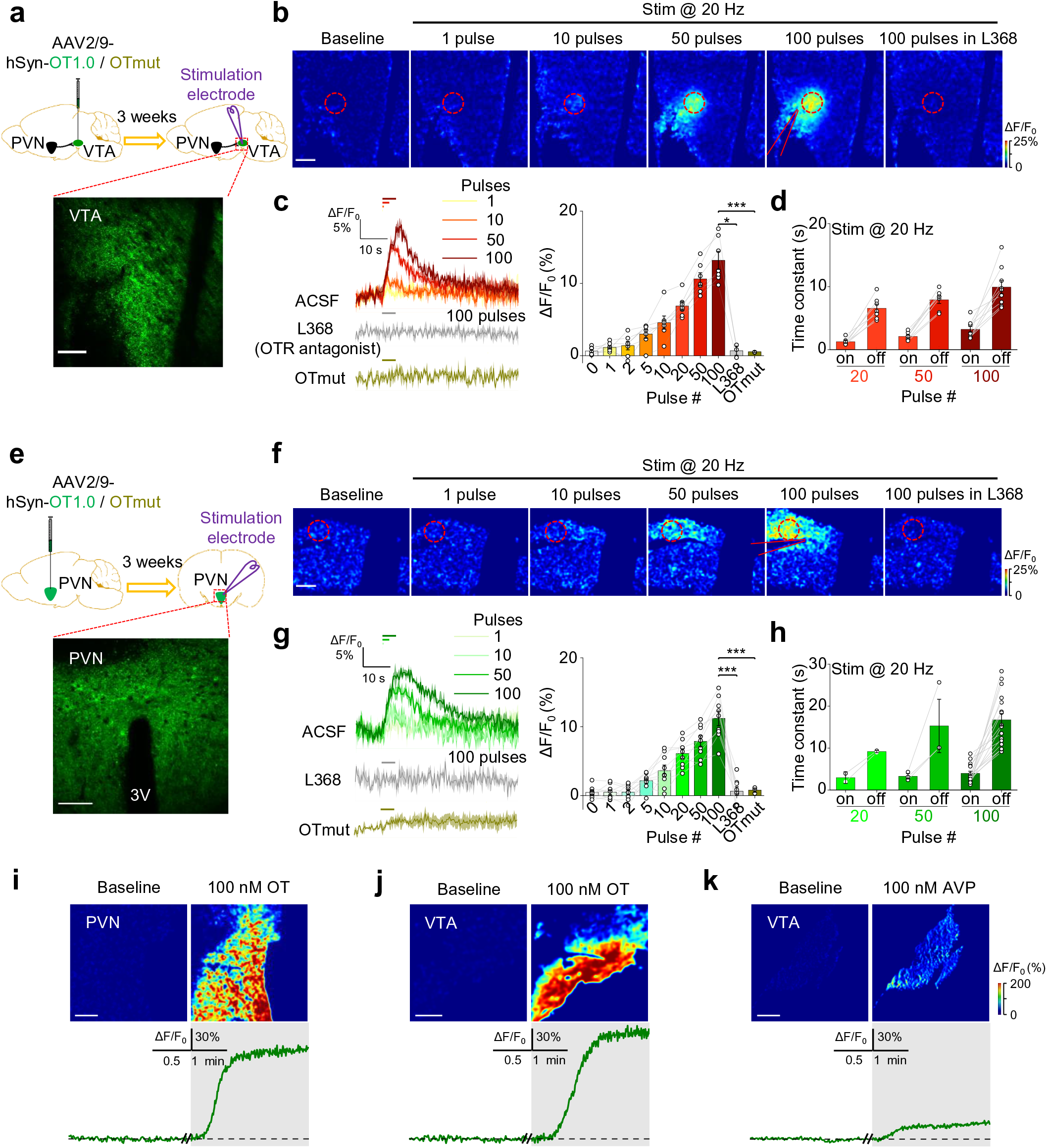
Activity-dependent axonal and somatodendritic OT release in brain slices. **a.** Top: schematic illustration depicting the experimental design for panels (**b-d**). Bottom: representative fluorescence image showing the expression of OT1.0 in the VTA. **b.** Example pseudocolor images of OT1.0-expressing brain slices at baseline and in response to 1, 10, 50, and 100 pulses electric stimuli (delivered at 20 Hz) and in response to 100 pulses in the presence of 5 μM L368. The dashed red circles indicate the ROI used to calculate the response, and the approximate location of the stimulating electrode is indicated. **c.** Representative traces (left) and summary (right) of the change in OT1.0 fluorescence in response to 1, 10, 50, and 100 pulses delivered at 20 Hz in ACSF, 100 pulses delivered in L368 (gray), and the change in OTmut fluorescence in response to 100 pulses. n=7 slices from 5 mice [7/5], 3/2, and 3/1 mouse for ACSF, L368, and OTmut, respectively. **d.** Summary of the rise time and decay time constants (τ_on_ and τ_off_) of the electrically evoked fluorescence increase in OT1.0-expressing slices in response to 20, 50, and 100 pulses delivered at 20 Hz. n=7, 7, and 9 slices for the 20-, 50-, and 100-pulse groups, respectively. **e.** Top: schematic illustration depicting the experimental design for panels (**f-h)**. Bottom: representative fluorescence image showing the expression of OT1.0 in the PVN. **f-h**. Similar to panels (**b-d**). In **g**, n=9/5, 9/5, and 3/1 for ACSF, L368, and OTmut, respectively; in **h**, n=2, 3, and 17 slices for the 20-, 50-, and 100-pulse groups, respectively. **i-k**. Example pseudocolor images (top) and fluorescence traces (bottom) of OT1.0-expressing slices containing the PVN (**i**) or VTA (**j, k**) before and after application of 100 nM OT (**i, j**) or AVP (**k**). \**p*<0.05, \*\*\**p*<0.001, and n.s., not significant. All scale bars represent 100 μm.

To measure somatodendritic OT release, we expressed OT1.0 under the control of hSyn promotor in the PVN and observed robust electrical stimulation‒induced responses, which had the rise time and decay time constants (τ_on_ and τ_off_) of 3.0-4.0 s and 9.2-16.7 s, respectively, and were blocked by L368 (**Fig. 2e-h**). Such responses were absent in slices expressing OTmut (**Fig. 2e, g**). In addition, bath application of 100 nM OT elicited a robust fluorescence increase in OT1.0 in both PVN-containing slices (**Fig. 2i**) and VTA-containing slices (**Fig. 2j**), with only slight response to 100 nM AVP (16% of peak ΔF/F_0_ compared to OT) measured in the VTA (**Fig. 2k**).

### Spatial and temporal analysis of axonal and somatodendritic OT release

Although the release of neuropeptides from large dense-core vesicles (LDCVs) is believed to occur on a much slower time scale than the release of classic neurotransmitters from synaptic vesicles (SVs)^41^, their precise spatial and temporal dynamics remain unclear. Therefore, we next compared the spatial and temporal kinetics of the axonal and somatodendritic OT release as well as glutamate (Glu) release, using OT1.0 sensor and iGluSnFR^42^, a genetically encoded glutamate sensor, respectively. Applying a standard train of electrical stimulation to acute brain slices (measured in the VTA and PVN) elicited OT1.0 fluorescence response with slower kinetics (with τ_on_ of 3.2-4.0 s and τ_off_ of 9.9-16.7 s) compared to iGluSnFR response (measured in the PVN, with τ_on_ of 0.5 s and τ_off_ of 4.5 s) (**Fig. 3a, b, f**). We then quantified ΔF/F_0_ at various distances from the release center and at various time points after the onset of stimulation and found that OT elicited a longer-lasting signal and diffused over a longer distance compared to Glu (**Fig. 3c, d, g**). The apparent diffusion coefficients are approximately 5×10^3^ μm^2^/s for OT in both the VTA and PVN, compared to 25.1×10^3^ μm^2^/s for Glu in the PVN (**Fig. 3e, h**). Our observations are consistent with the absence of a specific, rapid mechanism for recycling and degrading OT at the synaptic cleft, indicating that OT released from the axonal and somatodendritic compartments can diffuse slowly in the extrasynaptic space, serving as a long-lasting neuromodulator.

**Fig. 3:**
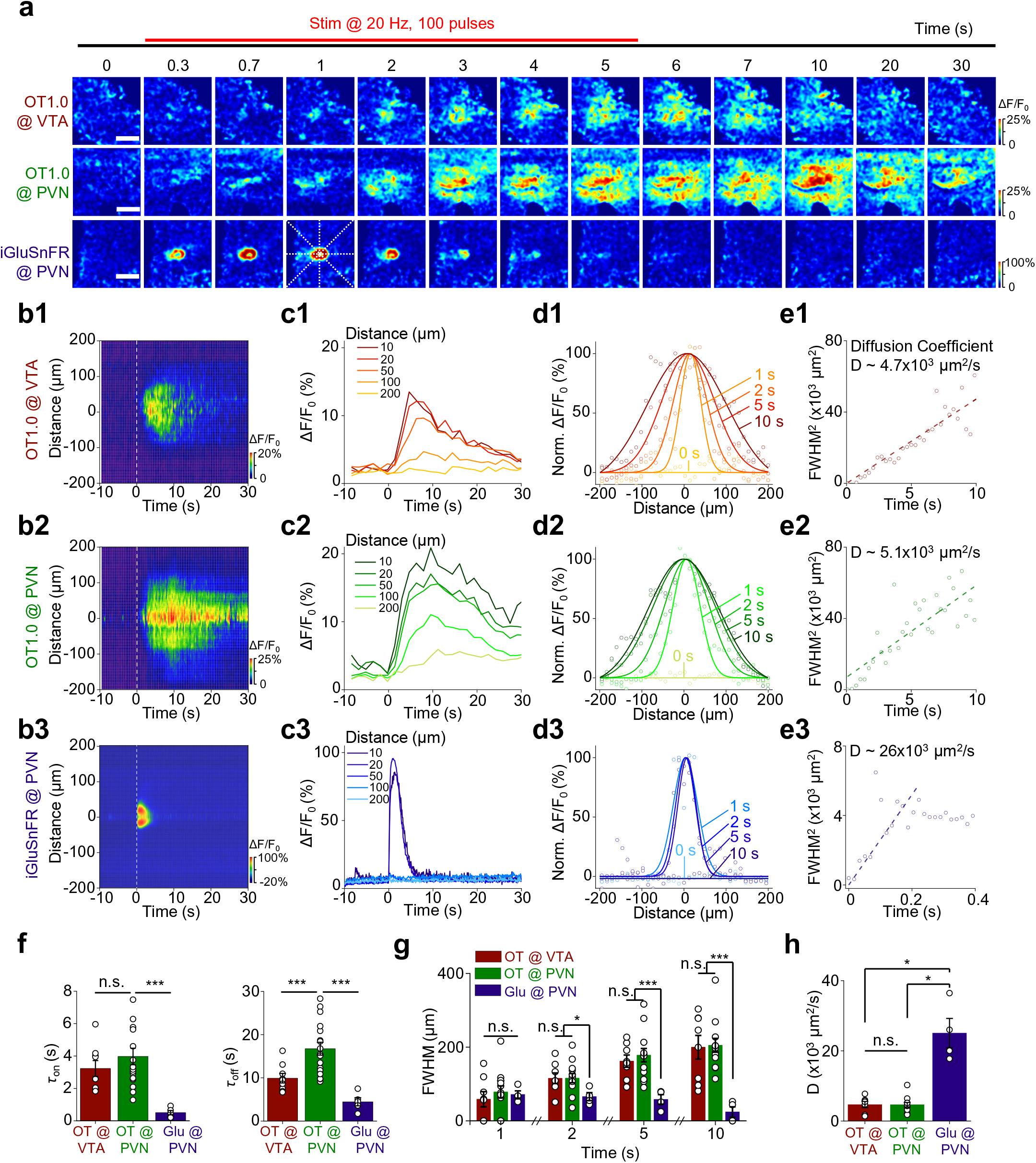
Probing the spatial and temporal dynamics of OT release in axonal and somatodendritic compartments. **a.** Example time-lapse pseudocolor images of OT1.0 expressed in the VTA (top row), OT1.0 expressed in the PVN (middle row), and iGluSnFR expressed in the PVN (bottom row) in acute brain slices. Where indicated, 100 electrical stimuli were applied at 20 Hz. The dashed lines were used to analyze the spatial and temporal dynamics. Similar results were observed for more than 6 slices for each group. Scale bars, 100 μm. **b.** Spatial profile of the evoked change in fluorescence shown in panel (**a**). The vertical dashed lines indicate the start of stimulation. Each heatmap shows the average of three trials conducted in one slice. **c.** Temporal dynamics of the data shown in **b** measured 10, 20, 50, 100, and 200 μm from the release center. The data were processed with 5x binning. **d.** Spatial dynamics of the data shown in **b** measured 1, 2, 5, and 10 s after the start of stimulation. Each curve was fitted with a Gaussian function. The data were processed with 5x binning and normalized to the peak response of each curve. **e.** Representative diffusion coefficients were measured by plotting the square of FWHM (full width at half maximum) against time based on the data shown in **d**. The diffusion coefficients were obtained by fitting a linear function using the FWHM^2^ calculated from the first 10 s (**e1**, **e2**) or the first 0.22 s (**e3**). **f.** Summary of τ_on_ and τ_off_ for the OT1.0 response in the VTA and PVN and iGluSnFR response in the VTA in response to 100 pulses delivered at 20 Hz (n=9, 17, and 4 slices, respectively). **g.** Summary of the diffusion distance (FWHM) of activity-dependent OT and Glu signals measured in **d** at the indicated time points; n=8 slices from 4 mice [8/4], 13/7, and 4/1 for OT @ VTA, OT @ PVN, and Glu @ PVN, respectively. **h.** Summary of the diffusion coefficients measured as in **e**; n=7/4, 12/6, and 4/1 mouse for OT @ VTA, OT @ PVN, and Glu @ PVN, respectively. \**p*<0.05, \*\*\**p*<0.001, and n.s., not significant.

### OT release employs compartment-specific calcium channels subtypes

Our finding that the activity-induced OT release had strikingly different kinetics compared to Glu release suggests that the release of these transmitters might be coupled to different release mechanisms such as different sources of Ca^2+^ influx. To test this hypothesis, we first measured OT and Glu signals in acute brain slices in response to the electrical stimulation in solutions containing different levels of Ca^2+^ (**Fig. 4a**). We found that both OT and Glu were released in a Ca^2+^-dependent manner, but with significantly different EC_50_ values (**Fig. 4a, b**). Specifically, OT release, particularly from the somatodendritic compartment, required higher concentrations of extracellular Ca^2+^. This observation is in line with high levels of neuronal activation and intracellular Ca^2+^ required for neuropeptide release^43^.

**Fig. 4:**
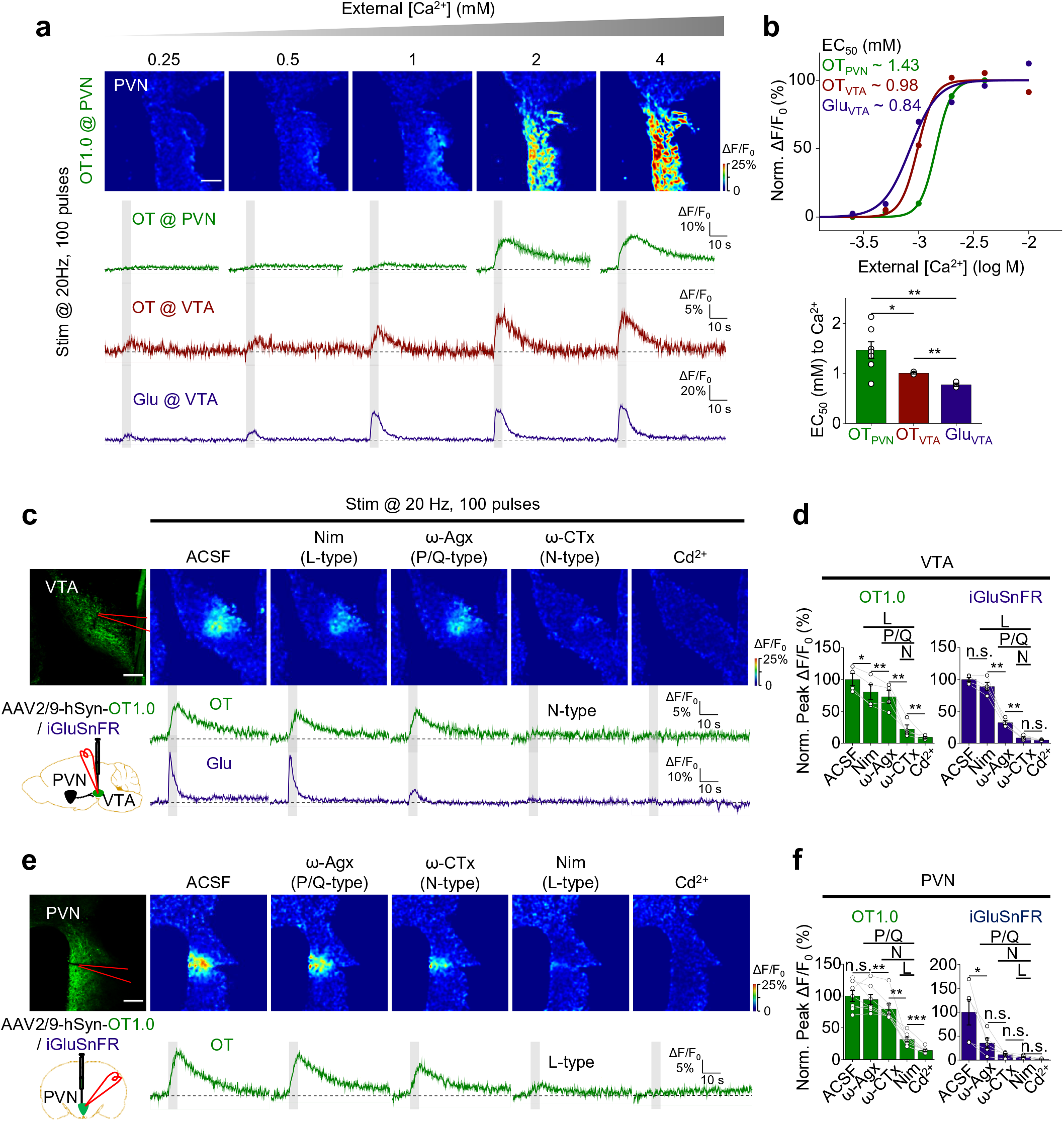
N-type and L-type voltage-gated Ca^2+^ channels support axonal and somatodendritic OT release, respectively. **a.** Pseudocolor two-photon images of OT release in the PVN (top row) and average ΔF/F_0_ traces (bottom) of OT1.0 in the PVN (green), OT1.0 in the VTA (red), and iGluSnFR in the VTA (blue) in response to 100 electrical stimuli applied at 20 Hz in the indicated concentrations of extracellular Ca^2+^. **b.** Summary of the normalized peak ΔF/F_0_ (top) measured in **a** at the indicated Ca^2+^ concentrations and summary of EC_50_ (bottom) values for OT1.0 in the PVN and VTA and iGluSnFR in the VTA; the data in the upper panel were normalized to the peak response measured in 4 mM Ca^2+^. n=9 slices from 4 mice [3/2], 3/2 mice, and 3/1 for OT @ PVN, OT @ VTA, and Glu @ VTA, respectively. **c.** Representative fluorescence image of OT1.0 (top left) and schematic drawing depicting the experimental strategy (bottom left). Example pseudocolor images (top) and traces (bottom) of the change in OT1.0 (green) and iGluSnFR (blue) fluorescence in response to 100 electrical stimuli delivered at 20 Hz in ACSF, the L-type VGCC blocker nimodipine (10 μM), the P/Q-type VGCC blocker ω-Agx-IVA (0.3 μM), the N-type VGCC blocker ω-CTx-GVIA (1 μM), or 200 μM Cd^2+^ to block all VGCCs (in this example, the same slice was sequentially perfused with the indicated blockers). **d.** Normalized peak responses for the data measured as shown in **c**; n=4 slices from 2 mice for OT1.0 and n=4 slices from 2 mice for iGluSnFR. **e, f**. Same as **c** and **d**, respectively, for OT1.0 and iGluSnFR expressed in the PVN; n=7 slices from 3 mice for OT1.0 and n=5 slices from 3 mice for iGluSnFR. \**p*<0.05 and \*\**p*<0.01. All scale bars represent 100 μm.

Oxytocinergic neurons express several subtypes of voltage-gated Ca^2+^ channels (VGCCs), including P/Q-, N-, L- and R-type^44–48^. The VGCC subtypes that are engaged in the activity-dependent OT release in the axonal or somatodendritic compartment remain largely unknown. We therefore examined which VGCC subtypes mediate axonal and somatodendritic OT release by sequentially treating slices containing either the VTA or PVN with blockers of specific VGCC subtypes and monitoring stimulation-induced OT release (**Fig. 4c, e**). We first confirmed that presynaptic Glu release in the VTA and PVN was inhibited by blocking both P/Q-type and N-type VGCCs with ω-agatoxin IVA (ω-Agx) and ω-conotoxin GVIA (ω-CTx), respectively^49, 50^ (**Fig. 4c, d, f, Extended Data Fig. 4c**). Notably, we found that axonal OT release depends mainly on N-type VGCCs while somatodendritic OT release is supported by L-type VGCCs (**Fig. 4c-f, Extended Data Fig. 4a, b**). Based on this pharmacological dissection, we conclude that activity-induced release of OT is Ca^2+^-dependent and employs compartment-specific VGCC subtypes.

### Activity-dependent OT release requires the SNARE complex

In neurons, SNARE (soluble N-ethylmaleimide-sensitive factor attachment receptor) complexes are essential for the release of neurotransmitter containing SVs^51^ and neuropeptide containing LDCVs^52^. With respect to SV fusion, the classic SNARE complex has been studied in detail and it consists of VAMP2^53^ (vesicle-associated membrane protein 2, also known as synaptobrevin 2), SNAP25^54^ (synaptosome-associated protein, 25 kDa), and syntaxin-1^55^. In contrast, the SNARE proteins mediating OT-containing LDCV fusion, with respect to different neuronal compartments, have not been fully characterized.

To characterize the SNARE proteins that mediate OT release, we expressed the light chain of botulinum toxin serotype A (BoNT/A)^54^ in the PVN to specifically cleave SNAP25 and subsequently measured OT and Glu release with OT1.0 sensor and iGluSnFR, respectively, in either the VTA (**Fig. 5a**) or PVN (**Fig. 5b, c**). As expected, Glu release evoked by 100 electrical pulses delivered at 20 Hz was reduced in BoNT/A-expressing PVN slices (**Fig. 5c**). Similarly, the stimulation-induced OT release detected by the OT 1.0 sensor was also significantly reduced in both the VTA and PVN slices expressing BoNT/A (**Fig. 5a, b**), indicating that SNAP25 plays a critical role in both axonal and somatodendritic OT release.

**Fig. 5:**
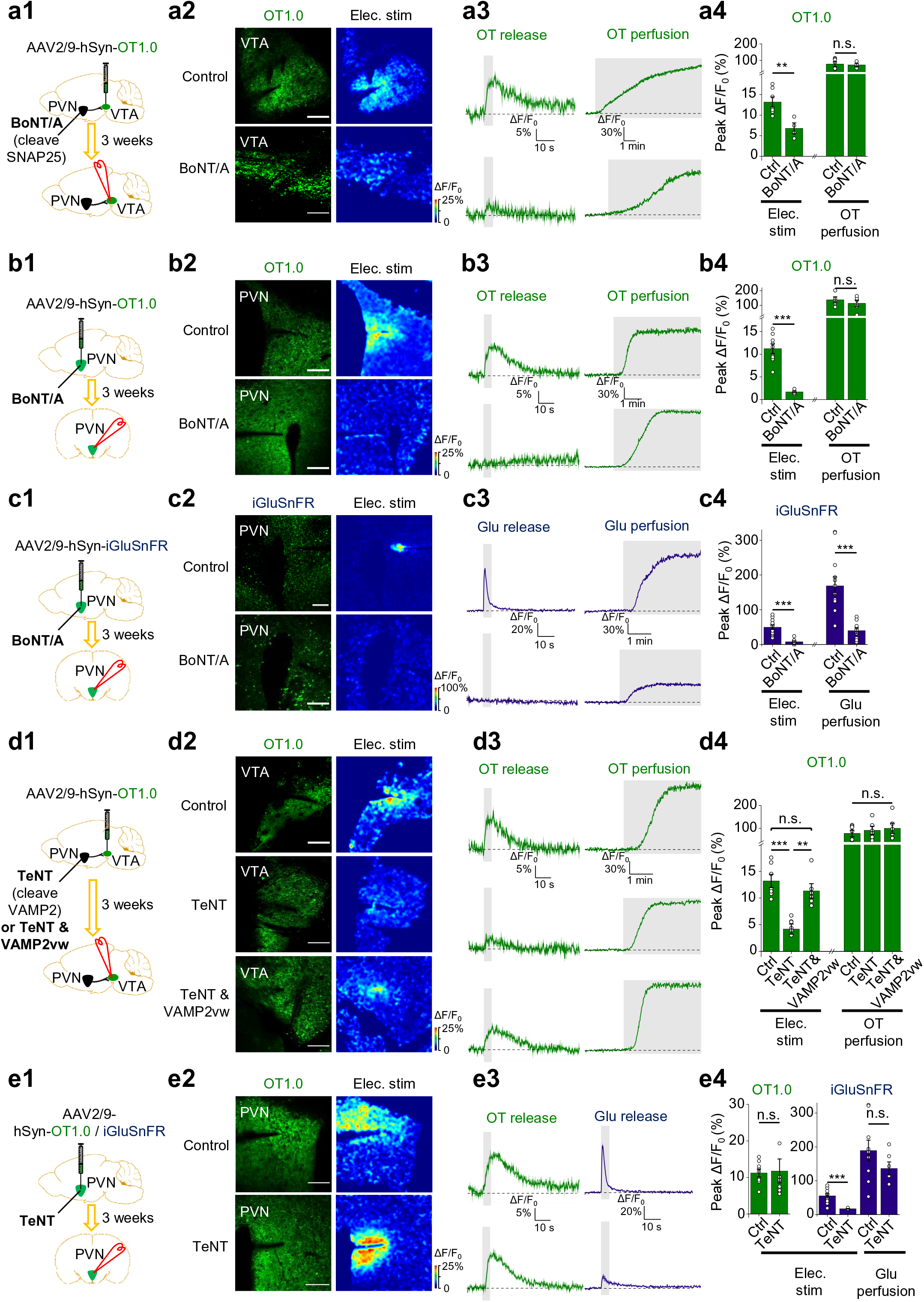
SNARE proteins play distinct roles in axonal and somatodendritic OT release. a-e. Schematic drawings depicting the experimental strategy (**a1-e1**), representative images showing the expression and peak response of OT1.0 sensor or iGluSnFR (**a2-e2**), average traces (**a3-e3**), and summary (**a4-e4**) of OT1.0 and/or iGluSnFR in response to 100 pulses stimulation delivered at 20 Hz or OT or Glu perfusion applied under the indicated conditions. BoNT/A, botulinum toxin serotype A; TeNT, tetanus toxin light chain. \*\**p*<0.01, \*\*\**p*<0.001, and n.s., not significant. All scale bars represent 100 μm.

Next, we expressed the tetanus toxin light chain (TeNT) in the PVN to specifically cleave VAMP1/2/3^53, 56^ and study its effect on the OT release in the VTA (**Fig. 5d**) and PVN (**Fig. 5e**) in a similar set of experiments. We found that TeNT expression significantly reduced axonal OT release in the VTA, and the reduction was rescued by co-expressing a TeNT-insensitive VAMP2 (VAMP2vw) with TeNT (**Fig. 5d**). Notably, somatodendritic OT release was unaffected by TeNT expression in the PVN, even though Glu release was significantly reduced (**Fig. 5e**), suggesting that VAMP2 does not serve as the principal VAMP protein in LDCV-mediated somatodendritic OT release. Taken together, these results indicate that the activity-dependent OT release requires the SNARE complex, with SNAP25 involved in both axonal and somatodendritic OT release, whereas VAMP2 essential only for axonal OT release.

### Optogenetic activation of OT neurons induces somatodendritic and axonal OT release *in vivo*

To address whether OT1.0 sensor can be used to detect changes in the OT level *in vivo*, we virally expressed OT1.0 sensor in the bed nucleus of stria terminalis (BNST) and measured its fluorescent response to rising concentrations of OT injected intraventricularly (**Fig. 6a**). A clear dose-dependent increase in fluorescence was observed over a 1,000-fold concentration range, with the maximum response reaching over 100% ΔF/F_0_ (**Fig. 6b, c**). The increase in OT1.0 sensor fluorescence was shown to be dependent on OT binding as it was blocked by pre-injection of oxytocin receptor antagonist, Atosiban (**Fig. 6d**). Control animals expressing OTmut in the BNST showed no response to OT injection at any dosage (**Fig. 6b, c**). We further examined the OT1.0 responses to AVP. Consistent with *in vitro* results and the natural selectivity of OTR, AVP-induced OT1.0 response was significantly lower than that induced by OT (**Fig. 6e**). Similarly, OT1.0-expressed in the PVN of female rats also responded to intraventricularly injected OT (**Extended Data Fig. 5**), indicating that OT1.0 sensor could be used to study OT release across rodent species.

**Fig. 6:**
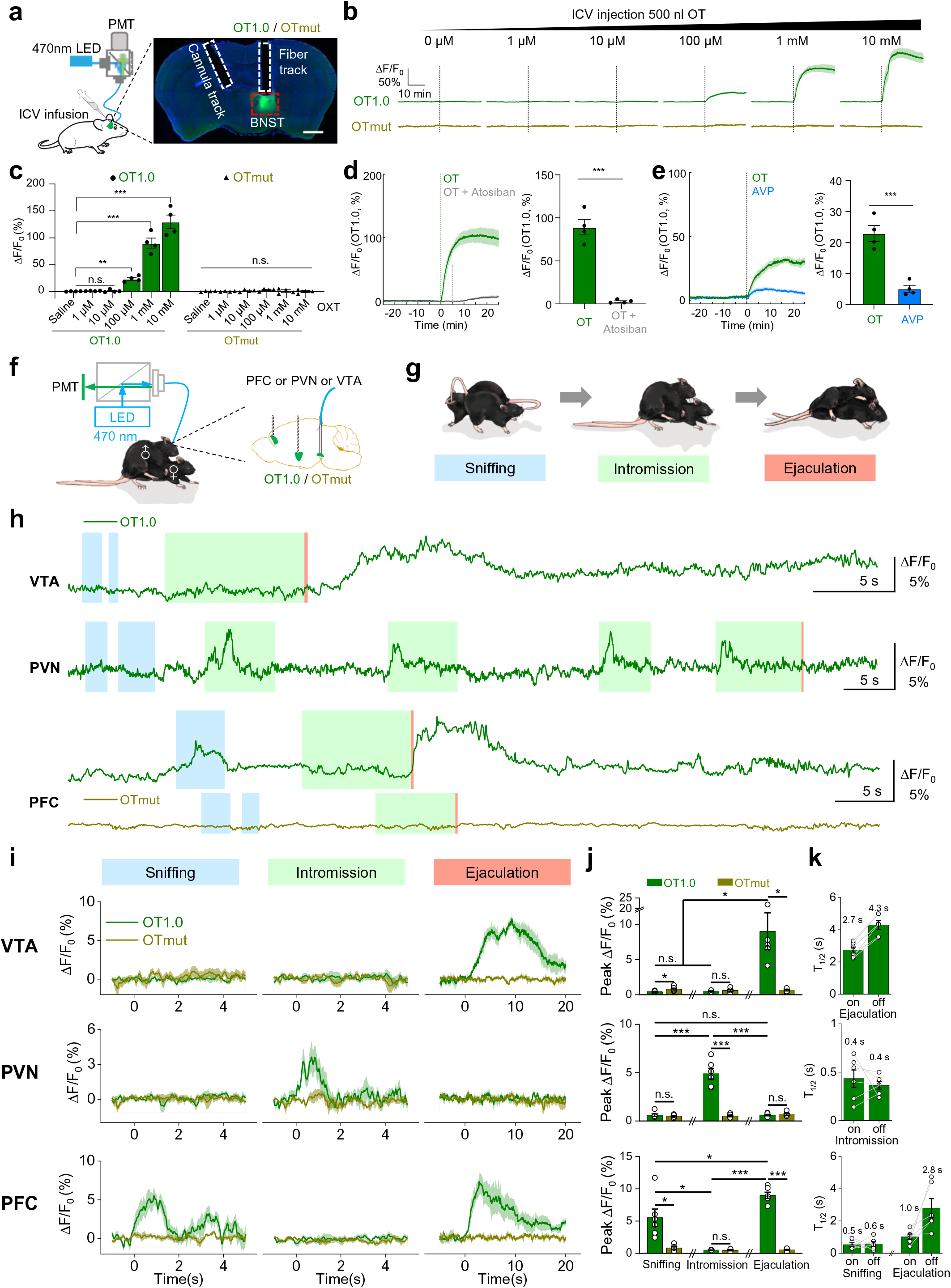
OT1.0 can be used to monitor OT release *in vivo* during male mating. **a**. Schematic illustrations depicting the *in vivo* ICV infusion experiments. An AAV expressing hSyn-OT1.0 or hSyn-OTmut was delivered into the BNST of male mice. After that, an optical fiber and a cannula were implanted in the BNST for monitoring fluorescence change during drugs infusion in freely moving mice. Scale bar, 1 mm. **b, c**. Representative traces (**b)** and summary (**c**) of the change in OT1.0 or OTmut fluorescence in response to the indicated concentrations of OT in freely moving mice. **d.** Example traces (left) and summary (right) of the ΔF/F_0_ in OT1.0 expressing mice in response to the application of 500 nl, 1 mM OT with (green) or without (blue) 500 nl, 50 mM Atosiban. **e.** Representative traces (left) and summary (right) of the ΔF/F_0_ in OT1.0 expressing mice in response to the application of 500 nl, 100 μM OT (blue) or 500 nl, 100 μM AVP (magenta). **f.** Schematic diagram depicting the experimental strategy for *in vivo* recording of OT1.0 in mice. An AAV expressing hSyn-OT1.0 was injected into the VTA, PVN, or mPFC of male mice; optical fibers were implanted in the corresponding brain regions 3 weeks later, and 470-nm light was used to excite the OT1.0 sensor during mating. **g.** Cartoon illustration of the three principal behaviors (sniffing, intromission, and ejaculation) exhibited by male mice when presented with a receptive female mouse. **h-k**. OT1.0 or OTmut was expressed in the VTA (top), PVN (middle), or mPFC (bottom) and imaged during male mating. Shown are representative traces of a single recording during male, with the various behaviors indicated (**h**), average time-locked traces from 5 individual behaviors (**i**), and summary of the peak responses (**j**) and rise time and decay time constants (**k**) measured during the indicated mating behaviors; n=6 mice per group. \**p*<0.05, \*\**p*<0.01, \*\*\**p*<0.001, and n.s., not significant.

To test whether OT1.0 sensor can detect endogenously released OT *in vivo* in mice, we expressed DIO-ChrimsonR-tdTomato^57^ in oxytocinergic PVN neurons together with either OT1.0 or OTmut in the medial prefrontal cortex (mPFC) of OT-Cre mice. We then optogenetically stimulated ChrimsonR-expressing cells in the PVN and measured OT1.0 sensor signal in the mPFC (**Extended Data Fig. 6a**). We found that optogenetic stimulation of the oxytocinergic PVN neurons with increasing numbers of light pulses delivered at 20 Hz evoked a time-locked, progressive increase in OT1.0 signal in the mPFC, which was blocked by i.p. injection of the OTR antagonist (L368) 5 minutes prior to the stimulation (**Extended Data Fig. 6b, c**). No increase in signal was measured in mice expressing OTmut in the mPFC, even after 10 s of optogenetic PVN stimulation (**Extended Data Fig. 6b, c**), indicating that OT release induces a reliable increase of OT1.0 fluorescence. The rise time and decay time constants (T_1/2_) of observed OT-mediated OT1.0 signal were 1.1-2.2 s and 1.4-4.2 s, respectively (**Extended Data Fig. 6d**). Similarly, optogenetic activation of the oxytocinergic PVN somas (**Extended Data Fig. 7a**) or SON-projecting axons (**Extended Data Fig. 7e**) with increasing frequencies of light pulses elicited a progressive increase in OT1.0 signal in PVN (**Extended Data Fig. 7b, c**) or SON (**Extended Data Fig. 7f, g**). Such responses were absent in rats during excitation at an isosbestic control wavelength (**Extended Data Fig. 7b, c, f, g**). The time constants (T_1/2_) of observed somatodendritic OT-mediated OT1.0 signal (measured in PVN, with rise time of 1.1-3.1 s and decay time of 1.2-9.7 s) were similar to axonal OT (measured in SON, with rise time of 0.5-6.8 s and decay time of 1.7-11.1 s) (**Extended Data Fig. 7d, h**). Taken together, these results confirm that the OT1.0 sensor can be used to measure compartmental OT release *in vivo* with high sensitivity, good specificity and rapid kinetics.

### Compartmental OT release during male mating behaviors in freely moving mice

OT has been reported to play a role in male mating behaviors^58–64^. It was shown to increase at PVN, VTA, medial preoptic area of the hypothalamus (mPOA), spinal cord, and other regions during male sexual behaviors to presumably regulate them via the OTR^64–67^. To study the dynamics of OT release during male mating behaviors, we performed fiber photometry recording of OT1.0 fluorescence in the VTA, PVN, and mPFC and observed region-specific responses during distinct phases of the behavior (**Fig. 6f, g**). In the VTA, OT1.0 increased (with rise time and decay time constants (T_1/2_) of 2.7 s and 4.3 s, respectively) during the ejaculation stage, but not during sniffing or intromission (**Fig. 6h-k, Supplementary Video 1**). In contrast, OT1.0 signal in the PVN increased during intromission (with rise time and decay time constants (T_1/2_) of 0.4 s and 0.4 s, respectively), but not during sniffing or ejaculation (**Fig. 6h-k, Supplementary Video 2**). Finally, OT1.0 signal in the mPFC was detected during sniffing (with rise time and decay time constants (T_1/2_) of 0.5 s and 0.6 s, respectively) and during ejaculation (with rise time and decay time constants (T_1/2_) of 1.0 s and 2.8 s, respectively) (**Fig. 6h-k, Supplementary Video 3**). As a negative control, no change in fluorescence was observed during any of the mating behaviors in the VTA, PVN, or mPFC of mice expressing OTmut (**Fig. 6f, h-j**), confirming that the signals measured in mice expressing OT1.0 were not movement artifacts. These results indicate that OT is transiently released in specific brain regions and from distinct neuronal compartments during specific mating stages, thereby encoding key information for specific features of sexual behavior.

## Discussion

Here, we report the development and characterization of a genetically encoded fluorescence indicator for monitoring extracellular OT both *in vitro* and *in vivo*. Our OT1.0 sensor has a wide dynamic range, good sensitivity and selectivity for OT over other neurotransmitters and neuropeptides, high spatial and temporal resolution, and negligible downstream coupling. In acute mouse brain slices, we show that OT1.0 can probe the release of OT from specific neuronal compartments, i.e., axonal OT release in the VTA and somatodendritic OT release in the PVN. Using this tool, we reveal differential molecular mechanisms underlying OT release from axonal and somatodendritic neuronal compartments. Finally, in a series of fiber photometry-based experiments in freely moving male mice, we show differential OT release in discrete brain regions during specific stages of mating behaviors, as summarized in **Fig. 7**.

**Fig. 7:**
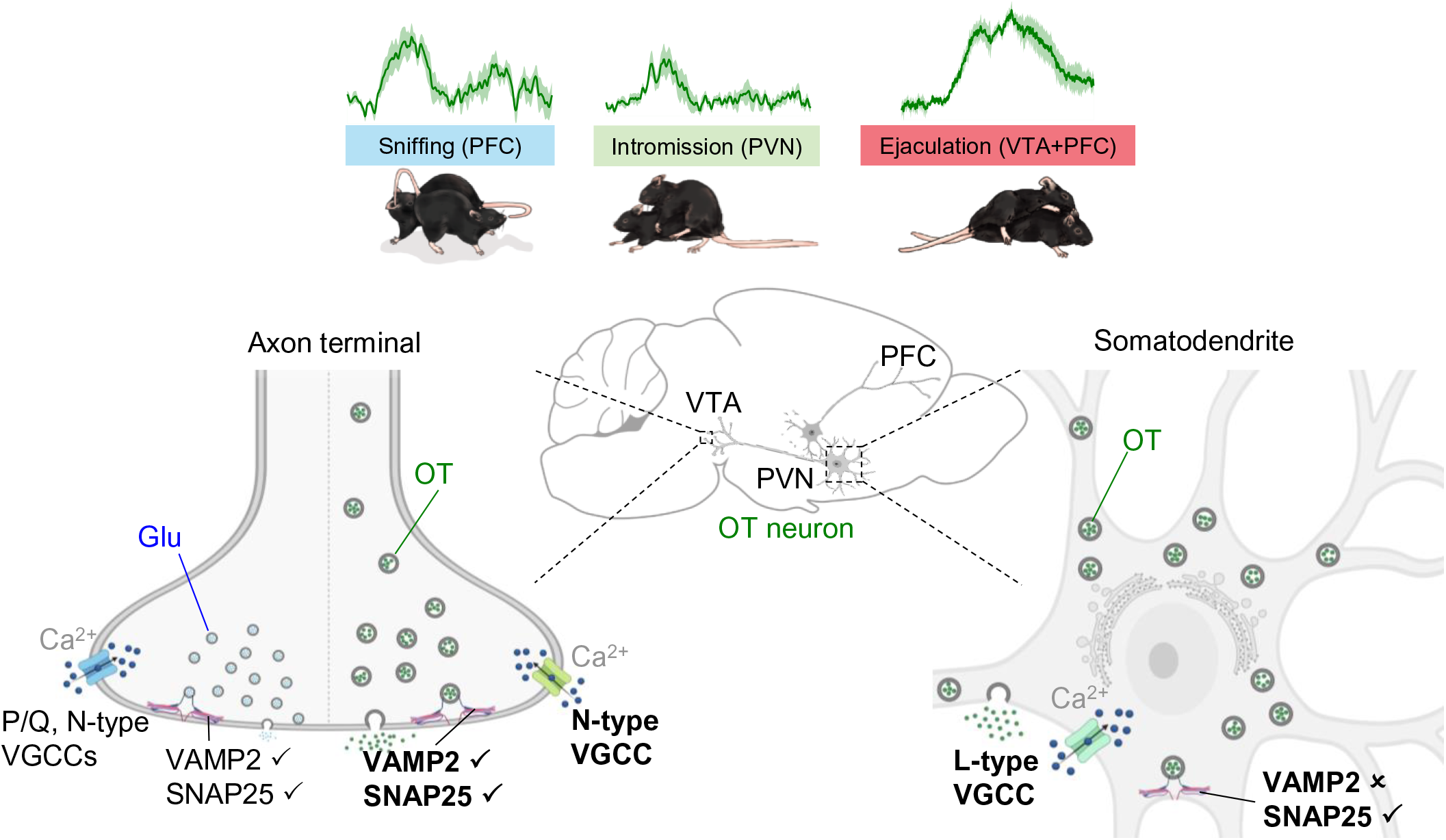
Model showing the molecular basis for axonal versus somatodendritic OT release. In the male brain during mating, OT is released from different compartments and via different mechanisms during the various behaviors. Top: during sniffing, OT is released primarily from the mPFC; during intromission, OT is released from the PVN; finally, during ejaculation OT is released from the VTA and mPFC. Bottom: axonal OT release is mediated primarily by N-type VGCCs and the SNARE proteins SNAP25 and VAMP2 (left). In contrast, somatodendritic OT release is mediated primarily by L-type VGCCs and SNAP25, but does not require VAPM2 (right). Note that the release of classic neurotransmitters such as glutamate (Glu) from small presynaptic vesicles is mediated by P/Q-type VGCCs, N-type VGCCs, SNAP25, and VAMP2.

Compared with other methods for monitoring OT release, our OT1.0 sensor has several distinct advantages. First, OT1.0 has a higher temporal resolution (τ_on_ ≈ 0.5 s) compared to microdialysis and OTR-iTango2, which are limited by a slow sampling rate (>5 min) and delayed reporter gene expression (∼48 h), respectively. Although fluorescent indicator protein‒tagged neuropeptides, such as NPY-pHluorin^68^, have been used to image LDCV-mediated peptide release, the relatively large fluorescent protein fused with the peptide renders precise spatiotemporal measurements of neuropeptide release difficult with this approach. Recently, Ino et al.^69^ reported a fluorescent OT sensor called MTRIA_OT_ based on the medaka OTR designed using a similar strategy as OT1.0. Compared to MTRIA_OT_, OT1.0 sensor has faster temporal dynamics, making it potentially more suitable for *in vivo* applications.

AVP, similarly to OT, can also be released from the somatodendritic compartment of the PVN neurons^70^. In the CNS, AVP and OT often display opposing roles, especially in the context of social anxiety and depression, and the balance between OT and AVP is important for maintaining a normal mental state^71^. OT1.0 has an approximately 12-fold higher apparent affinity for OT than AVP when expressed in cultured cells (**Extended Data Fig. 1c**), which is similar to the affinity of native mouse^72^, rat^73^, porcine^74^ and human^75^ OTRs. Further efforts could be applied to improve selectivity for OT over AVP, and vice versa, by modifying the ligand binding pockets of OT1.0. An OT indicator with even higher selectivity for OT over AVP is likely needed in order to help discriminate the specific functions of OT and AVP in the CNS.

Our *ex vivo* experiments showed that OT release from both the somatodendritic and axonal compartments is considerably slower than synaptic Glu release, consistent with faster kinetics of SVs fusion process compared to LDCVs^43^. Moreover, we showed that OT has a slower diffusion rate and diffuses over a longer distance than classic neurotransmitters such as Glu, which is possibly due to its relatively high molecular weight. Notably, we found that somatodendritic OT release has slower kinetics—in terms of both rise and decay times compared to axonal OT release. This might be due to the positive-feedback of OT onto the OT neurons. Indeed, OTR is expressed in the somas and dendrites of magnocellular oxytocinergic neurons^76^ where it couples to the Gq protein to increase intracellular Ca^2+^ concentration, thereby facilitating further OT release^77^. In addition, it is possibly caused by a slower replenishing of the more distant axonal release sites as compared to the somato-dendritic compartment^78^. Another possible mechanism that causes different kinetics of somatodendritic and axonal OT release is the involvement of different VGCCs and/or Ca^2+^ sensors. We found that somatodendritic OT release requires higher levels of extracellular Ca^2+^ than axonal release, suggesting that different Ca^2+^ sensors with different Ca^2+^ affinities are possibly present in these two compartments.

Moreover, we also showed here that different types of VGCCs predominantly mediate OT release from somatodendritic (L-type VGCCs) and axonal (N-type VGCCs) compartments; and that both SNAP25 and VAMP2 are required for the axonal OT release, while SNAP25—but not VAMP2—is required for somatodendritic OT release. The different regulation of VAMP2 partly provides the evidence for excluding the possible contribution of axonal OT release in PVN. Altogether, mentioned molecular differences can account for unique kinetics of axonal vs. somatodendritic OT release. While the transcriptome data reveal that P/Q-, N-type VGCCs, VAMP2 and SNAP25 are highly expressed in OT neurons^44, 46, 47, 79^, the expression patterns of these proteins are divergent between axonal and somatodendritic compartments of MNCs^45, 80^. With respect to VGCCs, electrophysiological study revealed that OT neurons are sensitive to L-type VGCCs blockage in cell body^81^, which is consistent with our results that L-type VGCCs mediate somatodendritic OT release. For the first time, here we showed that the N-type VGCCs are linked to axonal OT release in the VTA. Previously, R-type VGCCs were identified as a key regulator for OT secretion in posterior pituitary^82^. Taken together, this suggests existence of output specific release mechanisms in axonal projections of OT neurons. For SNARE proteins, the VAMP2 and SNAP25 were identified in axonal terminals, but were not enriched in soma or dendrite of OT neurons ^80^. Corresponding to this finding, our data suggest that other isoforms of synaptobrevin, not VAMP2, might be involved in somatodendritic OT release.

We also demonstrated that OT1.0 is an ideal tool to study OT dynamics *in vivo* and showed that OT is released during distinct phases of male sexual behaviors in the PVN containing cell bodies and dendrites of oxytocinergic neurons, as well as in the target regions of oxytocinergic neuron axons – the VTA and PFC. We observed that OT is specifically released in the PVN during intromission, in the PFC during sniffing and ejaculation, and in the VTA during ejaculation. In contradiction to our results, a microdialysis study revealed that OT level increases in the PVN after ejaculation^83^, which is possibly due to its poor spatial or temporal resolution. This mating-stage specific OT release in distinct brain regions is likely a consequence of differential control of axonal or somatodendritic oxytocin release^6^. We speculate that compartmental OT release is a key mechanism underlying temporally and spatially precise OT actions during behaviors. In line with this hypothesis, it was reported that somatodendritic OT release mediates bursting of OT neurons in lactating rats^21, 84^, providing a plausible mechanism by which somatodendritic OT release rapidly increases axonal OT secretion in posterior pituitary.

Given that the PVN contains both magnocellular and parvocellular oxytocinergic neurons, with specific electrophysiological properties, sizes, transcriptomes, and innervation patterns^40, 44, 85–89^, it will be crucial to determine whether the compartmentalized OT release occurs in all oxytocinergic neurons or specific subpopulations. We speculate that somatodendritic OT release mainly occurs in magnocellular OT neurons which have larger cell number and produce more OT peptide than parvocellular OT neurons in the PVN^6^, while the neuronal subtype responsible for axonal OT release in the VTA of mice remains controversial^39, 40^.

It is likely that compartmentalized release is displayed by other neurochemicals, such as dynorphin and brain-derived neurotrophic factor (BDNF)^4, 90^. The GRAB strategy proposed here can also be employed to develop new sensors for other neuropeptides that bind to G protein‒coupled receptors and answer fundamental physiological questions on diversity vs. similarity of peptide release mechanisms under diverse homeostatic or/and behavioral challenges occurring in various mammalian species, including primates. Furthermore, this strategy will be highly useful to disentangle the efficiency of peripherally administered OT (and other peptides) to pass through the blood brain barrier^91^, and subsequently affect human behavior^92, 93^, mitigating symptoms of mental diseases^94, 95^. In conclusion, our OT1.0 sensor is a robust new tool for studying OT functions under both physiological and pathophysiological conditions, and our finding that OT released from axonal and somatodendritic compartments is controlled by distinct and compartmentalized molecular mechanisms provides new insights into the complex actions of neuropeptides.

## Acknowledgements

This research was supported by the Beijing Municipal Science & Technology Commission (Z181100001318002 and Z181100001518004); grants from the National Natural Science Foundation of China (31925017, 31871087 and 81821092); grants from the NIH BRAIN Initiative (1U01NS113358 and 1U01NS120824), the Shenzhen-Hong Kong Institute of Brain Science (NYKFKT2019013); the Feng Foundation of Biomedical Research; and grants from the Peking-Tsinghua Center for Life Sciences and the State Key Laboratory of Membrane Biology at Peking University School of Life Sciences to Y.L.; and grants from the German Research Foundation (DFG) grants GR 3619/13-1, GR 3619/15-1, GR 3619/16-1, and the SFB Consortium 1158-2 to V.G. We thank Yi Rao for sharing the two-photon microscope, Xiaoguang Lei at PKU-CLS and the National Center for Protein Sciences at Peking University for providing support for the Opera Phenix high-content screening system. We thank Lara for packaging viral vectors used for in vivo experiments of A.K. and V.G. We thank Xiang Yu at Peking University, Bo Li at Cold Spring Harbor Laboratory, David Anderson at California Institute of Technology, Mark Andermann at Harvard University, Peter Kalugin at Harvard University and members of the Li lab for helpful suggestions and comments on the manuscript.

## Author contributions

Y.L. supervised the project. H.W. and L.G. performed the experiments related to the development, optimization, and characterization of the sensors in cultured cells. T.Q performed the two-photon imaging of OT dynamics in acute brain slices. L.M., T.O. and A.K. performed the *in vivo* ICV infusion experiments in mice and rats under the supervision of D.L. and V.G. P.W. performed the *in vivo* fiber photometry recording and optogenetic experiments in mice under the supervision of M.L. Y.T. performed the optogenetic experiments in rats under the supervision of R.S. All authors contributed to the interpretation and analysis of the data. T.Q. and Y.L. wrote the manuscript with input from all authors.

## Data and materials availability

The custom-written ImageJ macro, MATLAB code, and Arduino programs used in this study are available upon request.

## Methods

### Cell lines

HEK293T cells were obtained from American Type Culture Collection (ATCC). Cells expressing specific transgenes were selected using 2 μg/ml puromycin (Sigma). The HTLA cell line stably expressing a tTA-dependent luciferase reporter and the gene encoding the β-arrestin2-TEV fusion was a generous gift from B.L. Roth. All cell lines were cultured in DMEM (Gibco) supplemented with 10% (v/v) FBS (Gibco) and 1% penicillin–streptomycin (Gibco) at 37°C in 5% CO_2_.

### Primary neuronal cultures

Rat cortical neurons were cultured from postnatal day 0 (P0) Sprague-Dawley rat pups of both sexes (Beijing Vital River). Specifically, the brain was removed, the cortex was dissected out; the neurons were then dissociated in 0.25% trypsin-EDTA (Gibco), plated on 12-mm glass coverslips coated with poly-D-lysine (Sigma-Aldrich), and cultured in neurobasal medium (Gibco) containing 2% B-27 supplement, 1% GlutaMax (Gibco), and 1% penicillin-streptomycin (Gibco) at 37°C in 5% CO_2_.

### Animals

P0 Sprague-Dawley rats of both sexes (Beijing Vital River), male adult (P42-56) wild-type C57BL/6N (Beijing Vital River), male mice (Taconic) in SW background, female Sprague Dawley wild-type rats, male OT-IRES-Cre (Jackson Laboratory), and Ai14 mice were used to prepare primary neuronal cultures, acute brain slices and for *in vivo* experiment. Animals were housed at room temperature in 40-60% humidity under a 12-h/12-h light/dark cycle, with food and water available *ad libitum*. Procedures regarding animal experiments and maintenance were performed using protocols that were approved by the respective animal care and use committees at the Peking University, Chinese Institute for Brain Research, New York University and University of Heidelberg.

### Molecular cloning

DNA fragments were generated using PCR amplification with ∼25-bp primers (Tsingke). Plasmids were generated using the Gibson assembly method, and all plasmid sequences were verified using Sanger sequencing (Tsingke). All plasmids encoding the candidate OT sensors were cloned into the pDisplay vector (Invitrogen) with an upstream IgK leader sequence and a downstream IRES-mCherry-CAAX cassette for labeling the plasma membrane. cDNAs encoding the various OTRs were amplified from the human GPCR cDNA library, amplified from genomic DNA, or synthesized (Shanghai Generay Biotech), and the third intracellular loop (ICL_3_) of each OTR was swapped with the corresponding ICL_3_ in the GRAB_NE_ sensor. The plasmids for expressing the OT sensors in rat cortical neurons and mouse brain slices were cloned into the pAAV vector under the control of the human synapsin promoter (hSyn).

### Recombinant adeno-associated virus (AAV)

AAV2/9-hSyn-OT1.0 (1.80×10^13^ GC/ml, CIBR vector core or WZ Biosciences), AAV2/9-hSyn-OTmut (4.59×10^13^ GC/ml, WZ Biosciences), AAV2/9-hSyn-iGluSnFR (SF.iGluSnFR.A184V) (1.41×10^13^ GC/ml, WZ Biosciences), AAV2/9-EF1a-DIO-hM3Dq-mCherry (2.70×10^12^ GC/ml, BrainVTA), AAV2/9-CMV-BFP2-P2A-TeNT (1.54×10^14^ GC/ml, WZ Biosciences), AAV2/9-hSyn-jRGECO1a-P2A-VAMP2vw (2.02×10^13^ GC/ml, He Yuan Bioengineering), AAV2/9-hSyn-BFP2-P2A-BoNT/A (3.26×10^13^ GC/ml, He Yuan Bioengineering), and AAV2/9-CAG-DIO-ChrimsonR-tdTomato (1.30×10^13^ GC/ml, Shanghai Taitool Bioscience) were used to infect cultured neurons or were injected into specific mouse brain regions.

### Fluorescence imaging of cultured cells

An inverted confocal microscope (Nikon) equipped with a 40x/1.35-NA oil-immersion objective, a 488-nm laser, and a 561-nm laser was used for imaging; the GFP and RFP signals were collected using a 525/50-nm and 595/50-nm emission filter, respectively. Cultured cells expressing OT1.0 or OTmut were either bathed or perfused with Tyrode’s solution containing (in mM): 150 NaCl, 4 KCl, 2 MgCl_2_, 2 CaCl_2_, 10 HEPES, and 10 glucose (pH 7.4); where indicated, drugs and other compounds were delivered via a custom-made perfusion system or via bath application. An Opera Phenix high-content screening system (PerkinElmer) equipped with a 40x/1.1-NA water-immersion objective, a 488-nm laser, and a 561-nm laser was also used for imaging; the GFP and RFP signals were collected using a 525/50-nm and 600/30-nm emission filter, respectively. For imaging, fluorescence signals of the candidate OT sensors were calibrated using the GFP: RFP fluorescence ratio. To measure the response kinetics of the OT1.0 sensor, the line-scanning mode of the confocal microscope was used to record the rapid changes in fluorescence; a glass pipette containing 10 µM OT was placed near the surface of HEK293T cells expressing the OT1.0 sensor, and the OT was puffed onto the cell to measure τ_on_. To measure the dose-response curves for OT1.0 sensor shown in **Fig. 1g** and **Extended Data Fig. 1c, d**, solutions containing OT (ANASPEC or KS-V Peptide) or AVP (ANASPEC or KS-V Peptide), vasotocin (MedChemExpress), isotocin (MedChemExpress), inotocin (MedChemExpress) or nematocin (KS-V Peptide) with various concentrations were bath applicated in OT1.0 or OTmut expressed HEK293T cells and fluorescence was measured using an Opera Phenix high-content screening system (PerkinElmer).

### Tango GPCR assay

The wild-type bovine OTR (bOTR), OT1.0, or both was transfected into cell lines stably expressing a tTA-dependent luciferase reporter and the gene encoding the β-arrestin2-TEV fusion protein. After transfection, the cells were bathed in culture medium supplemented with various concentrations of OT, and then cultured for 12 h to allow the expression of luciferase. The culture medium was then replaced with Bright-Glo reagent (Fluc Luciferase Assay System, Promega) at a final concentration of 5 μM, and luminescence was measured using a Victor X5 multilabel plate reader (PerkinElmer).

### Calcium imaging of cultured cells

The plasmid of bOTR-ires-BFP or OT1.0-ires-BFP was transfected into HEK293T cells. After transfection, the cells were loaded with Cal590 (3 μg/ml, AAT Bioquest) for 50 minutes and solutions containing OT with various concentrations (0, 0.01, 0.1, 1 and 10 nM) and 100 μM ATP were orderly perfused and the Ca^2+^ response was measured using a confocal microscope (Nikon).

### Preparation and fluorescence imaging of mouse acute brain slices

Wild-type C57BL/6N mice or OT-Cre mice were deeply anesthetized by an i.p. injection of Avertin (500 mg/kg, Sigma-Aldrich), and then placed in a stereotaxic frame for injection of AAVs using a micro-syringe pump (Nanoliter 2000 Injector, WPI). For the data shown in **Fig. 2a-d**, AAVs expressing EF1a-DIO-hM3Dq-mCherry and/or hsyn-OT1.0 was injected (300 nl per site) into the left PVN of OT-Cre mice or OT-Cre x Ai14 mice using the following coordinates: AP: −0.75 mm relative to Bregma, ML: −1.5 mm, and DV: −4.95 mm relative to Bregma, at a 15° angle. For the data shown in **Fig. 2e-l and Fig. 3-5**, the indicated AAVs were injected in C57BL/6N mice (300 nl per site) into either the PVN as described above or the VTA using the following coordinates: AP: −3.2 mm relative to Bregma, ML: −0.5 mm, and DV: −4.6 mm relative to Bregma.

Three weeks after viral injection, mice were again deeply anesthetized with an i.p. injection of Avertin, and transcranial perfusion was performed using cold oxygenated slicing buffer containing (in mM): 110 choline-Cl, 2.5 KCl, 1 NaH_2_PO_4_, 25 NaHCO_3_, 7 MgCl_2_, 25 glucose, 0.5 CaCl_2_, 1.3 Na ascorbate, and 0.6 Na pyruvate. The brains were then rapidly removed and immersed into the oxygenated slicing buffer, after which the cerebellum was trimmed using a razor blade. The brains were then glued to the cutting stage of a VT1200 vibratome (Leica) and sectioned into 300-μm thick coronal slices. Brain slices containing the PVN or VTA were incubated at 34°C for at least 40 min in the oxygen-saturated Ringer’s buffer containing (in mM): 125 NaCl, 2.5 KCl, 1 NaH_2_PO_4_, 25 NaHCO_3_, 1.3 MgCl_2_, 25 glucose, 2 CaCl_2_, 1.3 Na ascorbate, and 0.6 Na pyruvate. For two-photon imaging, the slices were transferred into an imaging chamber in an FV1000MPE (Olympus) or Bruker two-photon microscope equipped with a 25× /1.05-NA water-immersion objective and a mode-locked Mai Tai Ti:Sapphire laser (Spectra-Physics) tuned to 920 nm with a 495-540-nm filter for measuring fluorescence. For electrical stimulation, a homemade bipolar electrode (cat. #WE30031.0A3, MicroProbes) was placed onto the surface of the brain slice near the VTA or PVN under fluorescence guidance. Imaging and stimulation were synchronized using an Arduino board with a custom-written program. The experiments for probing the spatial and temporal kinetics of Glu release were recorded at a video frame rate of 0.0171 s/frame, with 256×256 pixels per frame. All other stimulation experiments were recorded at video frame rates of 0.3583 or 0.3259 s/frame, with 256×192 or 256×256 pixels per frame. The stimulation voltage was set at 5-8 V, and the duration of each stimulation was 1 ms. Where applicable, drugs and other compounds were applied to the imaging chamber by perfusion in ACSF at a flow rate of 4 ml/min.

### *In vivo* fiber photometry recording of OT1.0 responses during ICV infusion of drugs in mice

Adult WT male mice (Taconic) on the SW background were used for surgery. During surgery, mice were anesthetized with 1%-2% isoflurane and mounted onto a stereotaxic device (Kopf Instruments Model 1900). 500nl AAV9-hSyn-OT1.0 or AAV9-hSyn-OTmut viruses were delivered into the BNST (AP: −0.45 mm, ML: −0.9 mm, DV: −3.6 mm) through a glass capillary using nanoinjector (World Precision Instruments, Nanoliter 2000). After virus injection, a 400-µm optical fiber assembly (Thorlabs, FR400URT, CF440) was inserted 300 µm above the virus injection site and secured onto the skull using an adhesive dental cement (C&B Metabond, S380), at the same time a cannula was inserted into the right-side lateral ventricle and secured onto the skull using an adhesive dental cement (C&B Metabond, S380). Three weeks after surgery, mice were head-fixed on a running wheel and fluorescence signals of the sensor were acquired as described previously ^96^. Mice were recorded for 25 min and subsequently ICV infused with various amounts (time interval between 2 trials ≈ 24 h) of OT and AVP dissolved in 0.5 µl of saline and recorded for further 25 min. The OTR antagonist Atosiban (50 mM in 0.5 µl saline) was ICV injected 5 minutes prior to the OT infusion. The drug-induced responses were calculated as the mean fluorescence level after each drug injection minus the mean fluorescence level before drug injection.

### *In vivo* fiber photometry recording of OT1.0 responses during ICV infusion of OT in rats

Adult female Sprague Dawley rats were anesthetized with 2-4% isoflurane and mounted onto the stereotaxic frame (Kopf Instruments). 500 nL of AAV9-hSyn-OT1.0 viruses were delivered into the PVN (AP: −1.8 mm, ML: −0.35 mm, DV: −8 mm relative to Bregma) using a glass microinjection pipettes connected to a syringe pump. At least 2 weeks after the viral injection, rats were anesthetized again with isoflurane and mounted onto the stereotaxic frame. Optical fibers (400 µm) were placed above the PVN region (AP: −1.8 mm, ML: −2 mm, DV: −7.8 mm relative to Bregma, at an 14° angle), while glass microinjector pipettes connected to a syringe pump were lowered into the lateral ventricle (AP: −0.7 mm, ML: 1.8 mm, DV: −4 mm relative to Bregma). Fiber photometry experiment was performed using Bundle-imaging Fiber Photometry setup (Doric Lenses). OT1.0 was stimulated using a 400-410 nm (isosbestic) and 460-490 nm excitation LEDs; emitted 500-550 nm fluorescence was detected with a CMOS camera in interleaved acquisition mode. The signal was recorded for 3 minutes before the ICV OT injection (1 µL, 10 mM) and 20 minutes afterwards. To verify the optical fiber placement and OT1.0 expression, selected rats were transcardially perfused with 100 ml of saline followed by 100 ml of 4% formaldehyde in PBS. Postfixed brains were cut into coronal sections (50 µm) and analyzed with epifluorescent microscope (Nikon). Sections were stained against GFP to visualize the sensor expression.

### *In vivo* fiber photometry recording of OT1.0 responses during optogenetic activation in rats

Adult female Sprague Dawley rats were anesthetized with 2-4% isoflurane and mounted onto the stereotaxic frame. As described previously^97^, an AAV expressing oxytocin promoter (pOT) - ChrimsonR-tdTomato was injected into the PVN, and an AAV expressing hSyn-OT1.0 was injected into the PVN or SON of SD-rat using the following coordinates (PVN, AP: −1.8 mm, ML: 0.4 mm, DV: −8.0 mm relative to Bregma; SON, AP: −1.2 mm, ML: 1.8 mm, DV: −9.2 mm relative to Bregma); 3 weeks after virus injections, an optical fiber (400 um, 0.5NA) was implanted in the PVN or SON to activate oxytocin neuron somas or axons and record oxytocin release. To this purpose it was coupled with a fiber patch cord (200 um, 0.37NA) and linked to an integrated fluorescence minicube (Doric ilFMC5). The minicube was connected to a fluorescence detector head to record the 525 nm fluorescence emission signal and connected to a double wavelength LED (20 uW/mm^2^) to excite the OT1.0 sensor at isosbestic control wavelength (405 nm) or OT-sensitive sensor wavelength (465 nm). In addition, it was connected to a 593 nm LED (10 mW/mm^2^) to activate ChrimsonR through the same fiber core without bleedthrough into the 525 nm detection channel. Traces are the average response of 30 individual trials in the same rat, light colour indicates the standard deviation.

### *In vivo* fiber photometry recording of OT dynamics during optogenetic activation or sexual behavior

Male adult wild-type C57BL/6N (Beijing Vital River) or male adult OT-IRES-Cre (Jackson Laboratory) mice were deeply anesthetized with an i.p. injection of Avertin and then placed in a stereotaxic frame for AAV injection. For optogenetics, AAVs expressing hSyn-OT1.0 or hSyn-OTmut were injected (300 nl per site) into the left mPFC using the following coordinates: AP: +1.9 mm relative to Bregma, ML: −0.3 mm, and DV: −1.8 mm relative to Bregma; in addition, an AAV expressing CAG-DIO-ChrimsonR-tdTomato was injected (300 nl per site) into the left PVN of male OT-IRES-Cre mice using the following coordinates: AP: −0.75 mm relative to Bregma, ML: −1.5 mm, and DV: −4.95 mm relative to Bregma at a 15° angle. Mice were treated with an i.p. injection of either saline or L368 (Tocris, 10 mg/kg) 30 minutes before light stimulation. For the sexual behavior experiments involving male wild-type mice, AAVs expressing hSyn-OT1.0 or hSyn-OTmut were injected (300 nl per site) into the left VTA using the following coordinates: AP: −3.2 mm relative to Bregma, ML: −0.5 mm, and DV: −4.6 mm relative to Bregma, the left PVN using the following coordinates: AP: −0.75 mm relative to Bregma, ML: −1.5 mm, and DV: −4.95 mm relative to Bregma at a 15° angle, or the left mPFC using the following coordinates: AP: +1.9 mm relative to Bregma, ML: −0.3 mm, and DV: −1.8 mm relative to Bregma. Optical fibers (105-μm core/125-μm cladding) were implanted in the mPFC, VTA, and/or PVN 3 weeks after AAV injection. Fiber photometry was recorded in the mPFC, VTA, and/or PVN using a 470-nm laser at 50 μW for OT1.0 or OTmut, and ChrimsonR expressed in the PVN was stimulated using a 593-nm laser at 10 mW (10-ms pulses were applied at 20 Hz for 0.25-10 s). A 535/50-nm filter was used to collect the fluorescence signal from OT1.0 or OTmut. The animal’s sexual behaviors were recorded using the commercial video acquisition software StreamPix 5 (Norpix), and behaviors were annotated and tracked using custom-written MATLAB codes (MATLAB R2019a, MathWorks). After excluding mice with incorrect fiber placement, we analyzed 6 out of 18 mice, 6 out of 18 mice, and 6 out of 16 mice with significant fluorescence change during mating behavior in PVN, VTA and PFC, respectively.

### Data analysis

Imaging data obtained from cultured cells and acute brain slices were processed using ImageJ software (NIH) with a custom-written macro. The change in fluorescence (ΔF/F_0_) was calculated using the formula (F−F_0_)/F_0_, in which F_0_ is the baseline fluorescence signal. Summary data are presented as the mean±s.e.m. Statistical analyses were performed using GraphPad Prism 8. The two-tailed Student’s *t*-test or one-way ANOVA with Tukey’s multiple comparisons test were performed where appropriate, and differences with a *p*-value <0.05 were considered significant. The traces and summary graphs were generated using OriginPro 9.1 (OriginLab).

## Supplementary information

**Suppplementary Video 1.**
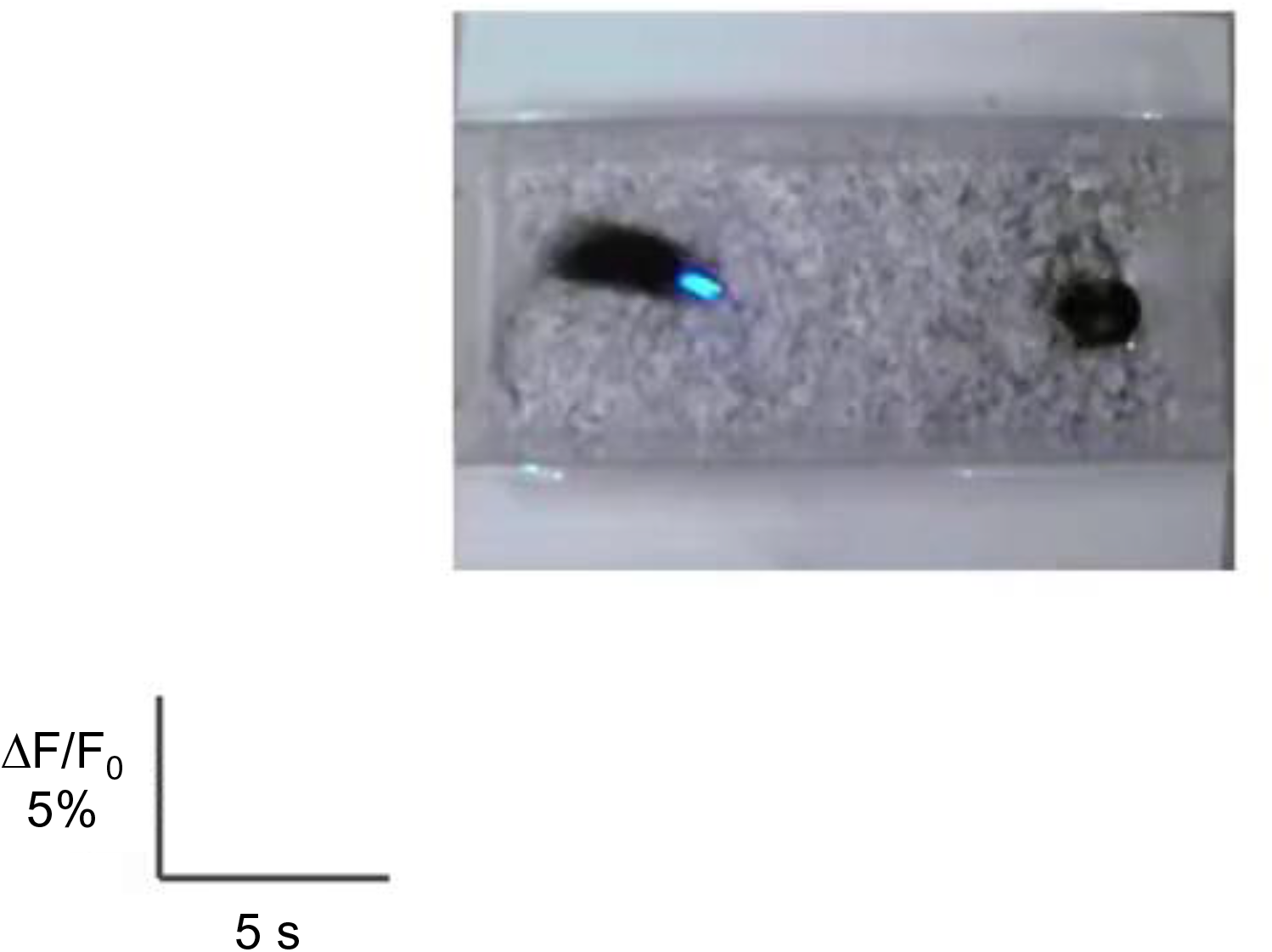
Single-trial measurement of OT1.0 fluorescence in the VTA in the male mouse brain during mating (related to Fig. 6).

**Supplementary Video 2.**
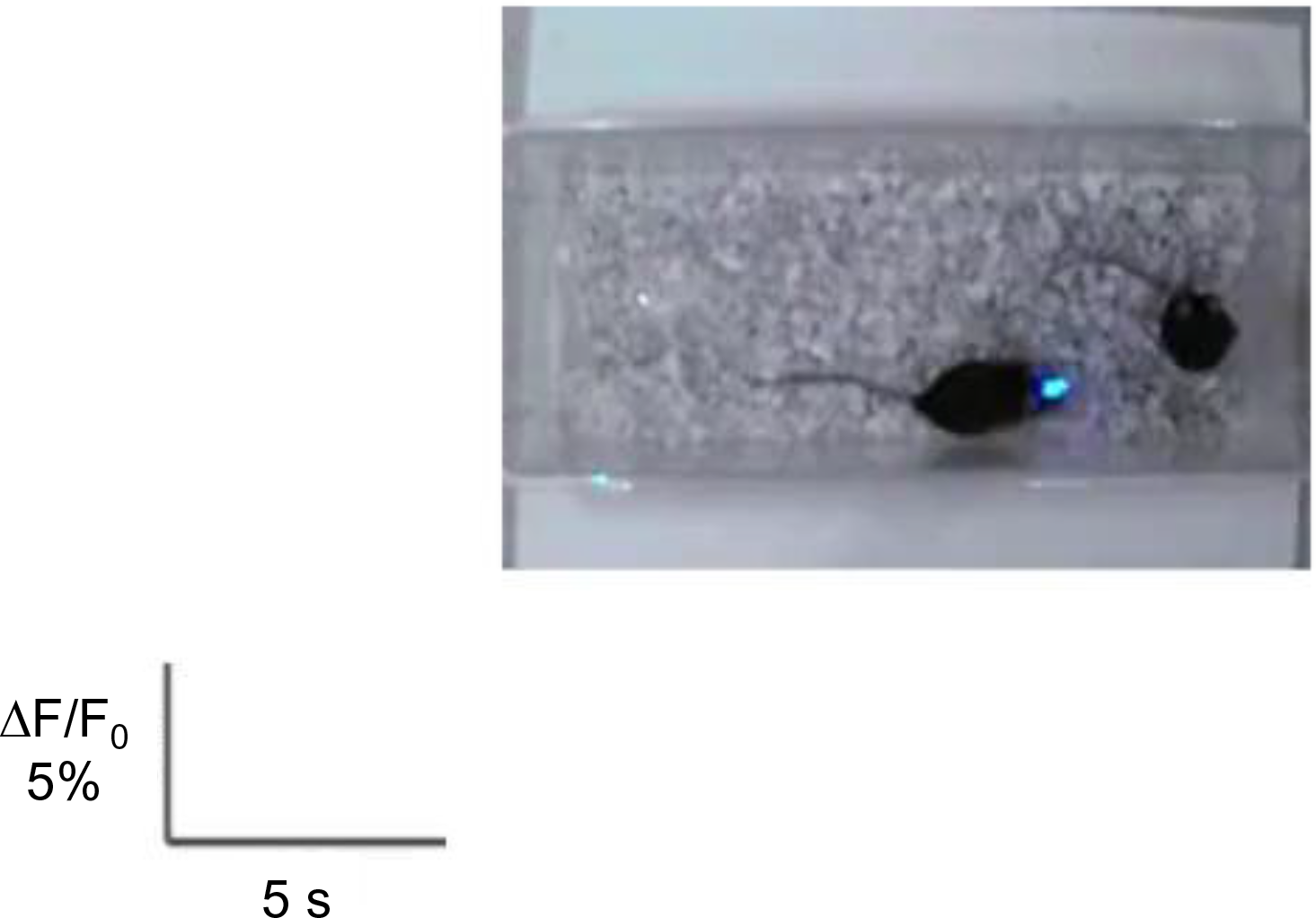
Single-trial measurement of OT1.0 fluorescence in the PVN in the male mouse brain during mating (related to Fig. 6).

**Supplementary Video 3.**
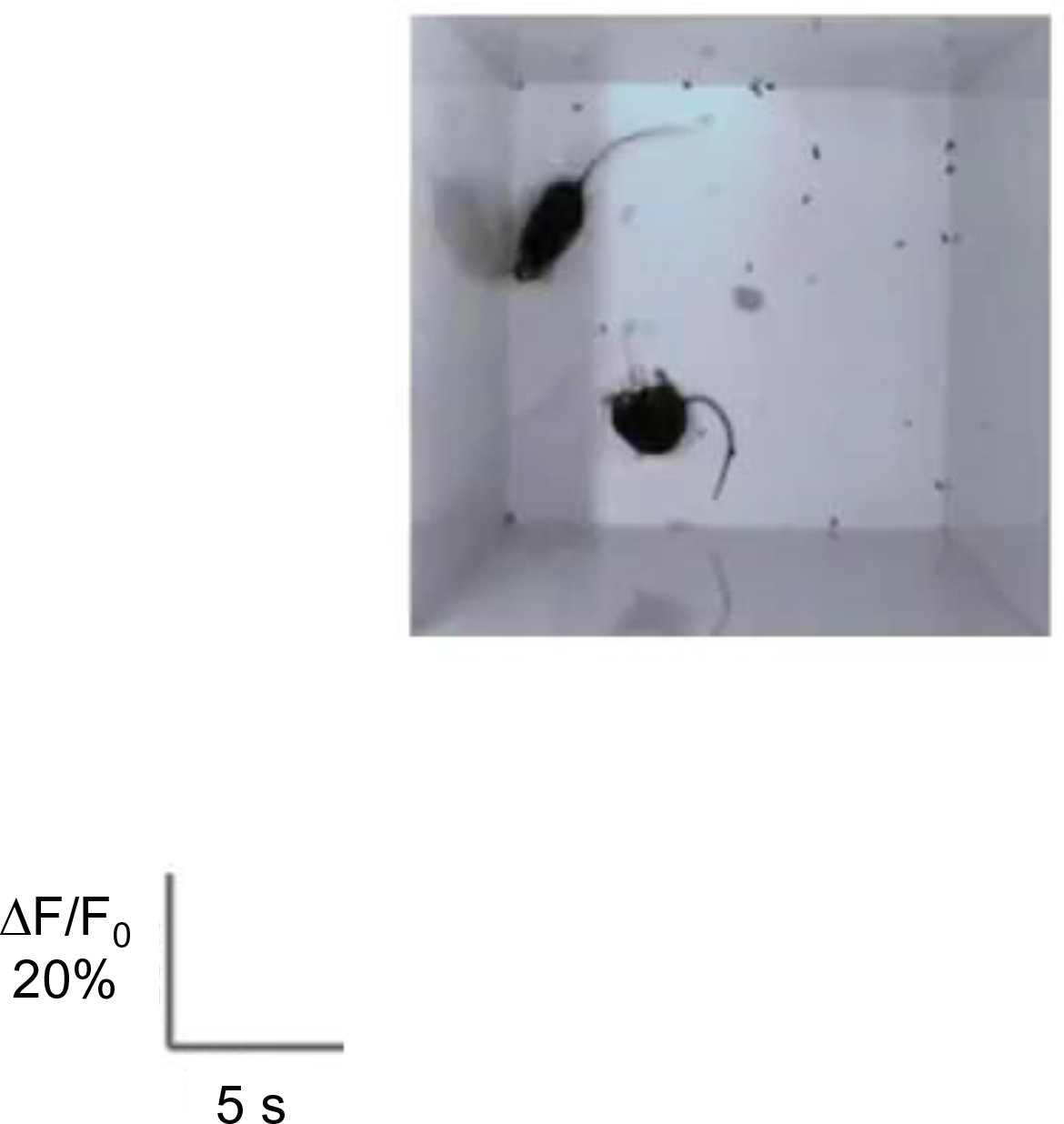
Single-trial measurement of OT1.0 fluorescence in the mPFC of the male mouse brain during mating (related to Fig. 6).

---

**Extended Data Fig. 1:**
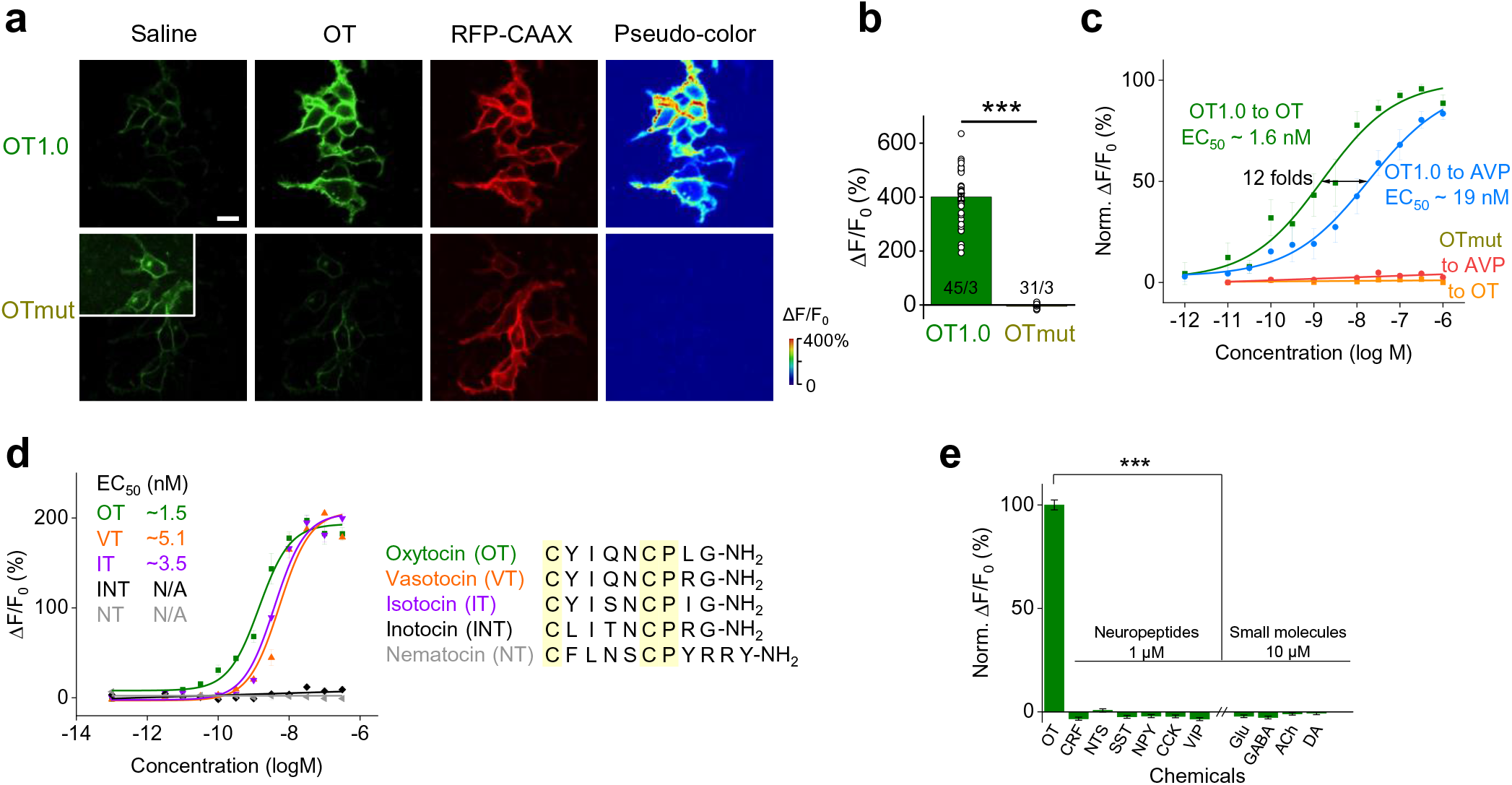
Characterization of GRAB_OT_ sensors in HEK293T cells. **a.** Representative images of OT1.0 and OTmut expressed in HEK293T cells in saline and in the presence of 100 nM OT. Also shown is RFP-CAAX expression, showing localization at the plasma membrane. The images at the right show the change in OT1.0 and OTmut fluorescence in response to OT application. White rectangle with enhanced contrast showing OTmut expressing HEK293T cells in saline. Scale bar, 20 μm. **b.** Summary of the peak change in OT1.0 and OTmut fluorescence measured in HEK293T cells in response to 100 nM OT. **c.** Dose–response curves for OT1.0 and OTmut expressed in HEK293T cells in response to the indicated concentrations of OT and AVP, with the corresponding EC_50_ values shown. The data were normalized to the maximal response measured in OT group. The dosage curves of OT1.0 to OT/AVP were averaged from 9 individual trials, with 3-4 wells per trial. **d.** Dose-response curves for OT1.0 expressed in HEK293T cells in response to the indicated concentrations of OT and its orthologous peptides, with amino acid sequence alignment shown. **e.** Summary of the peak change in OT1.0 fluorescence measured in HEK293T cells in response to the indicated compounds applied at 1 μM (CRF, NTS, NPY, and VIP) or 10 μM (Glu, GABA, Gly, DA, NE, and 5-HT), normalized to the peak response measured in OT; n=3 wells per group. CRF, corticotropin-releasing factor; NTS, neurotensin; NPY, neuropeptide Y; VIP, vasoactive intestinal peptide; Glu, glutamate; GABA, γ-aminobutyric acid; Gly, glycine; DA, dopamine; NE, norepinephrine; and 5-HT, 5-hydroxytryptamine (serotonin). \*\*\**p*<0.001.

**Extended Data Fig. 2:**
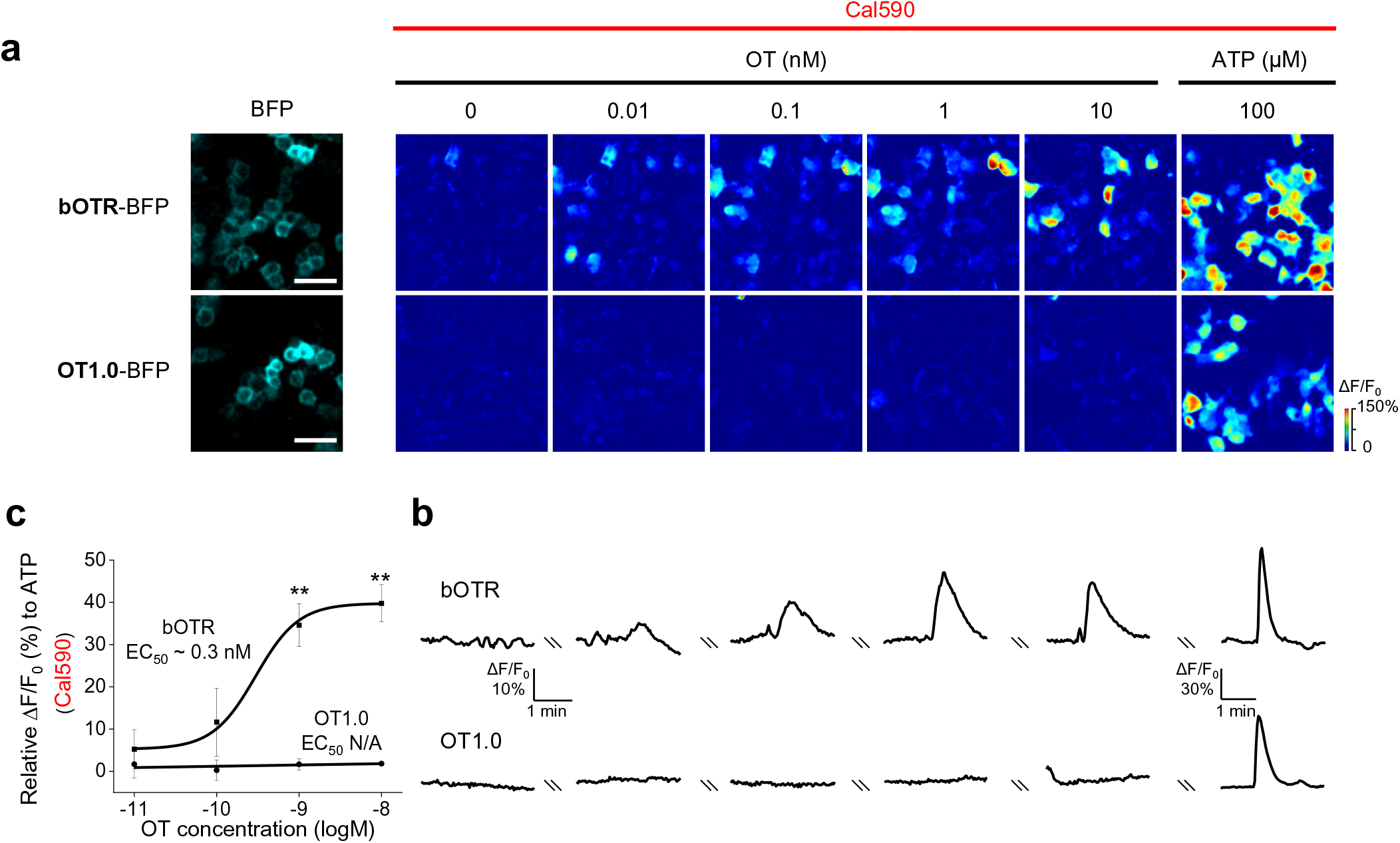
Negligible downstream Gq-dependent calcium signaling coupling of OT1.0 sensor in HEK293T cells. **a**, **b**. Representative expression images in BFP channel, pseudocolor images (top) and ΔF/F_0_ traces (bottom) showing the Ca^2+^ response to the indicated concentrations of OT or ATP in HEK293T cells expressing bOTR-BFP (**a**) or OT1.0-BFP (**b**). Scale bar, 50 μm. **c**. Summary of peak Ca^2+^ ΔF/F_0_ for bOTR or OT1.0 expressed HEK293T cells corresponding to (**a** and **b**) at indicated OT concentrations, with the corresponding EC_50_ value shown. The data were normalized to the peak response measured in 100 μM ATP. n=3 coverslips for each group. **p < 0.01 and n.s., not significant.

**Extended Data Fig. 3:**
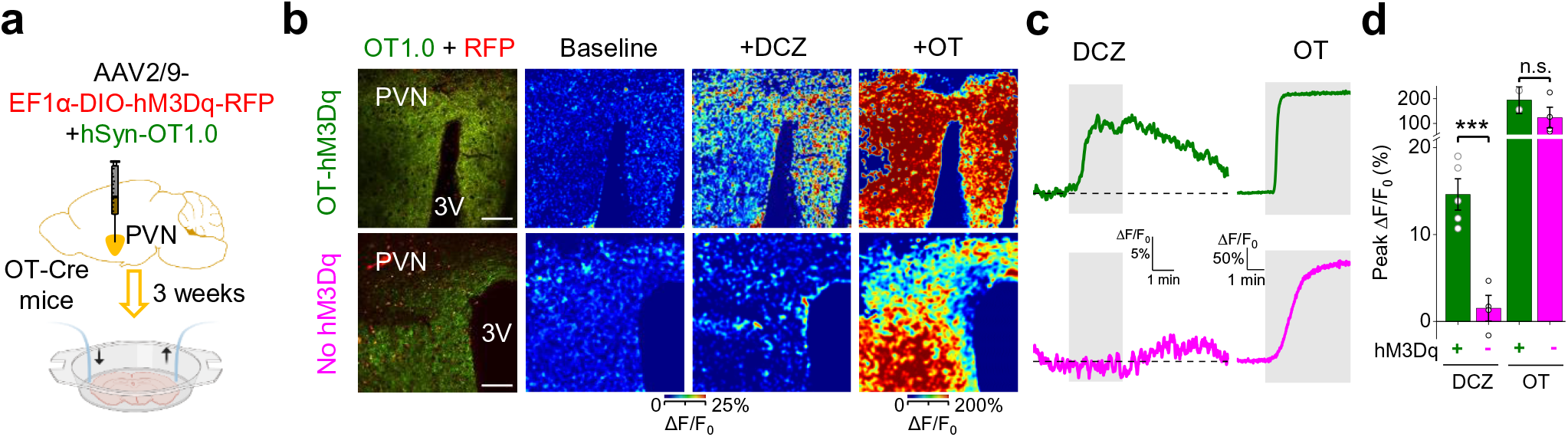
Chemogenetic activation of oxytocinergic neurons induces OT release. **a.** Schematic diagram depicting the chemogenetic activation experiments. A mixture of AAVs (EF1α-dio-hM3Dq-mCherry and hSyn-OT1.0) was injected into the PVN of OT-Cre mice. As a control, hSyn-OT1.0 was injected into the PVN of OT-Cre x Ai14 mice (no-hM3Dq). The PVN and third ventricle (3V) are indicated. **b.** Left: representative 2-photon microscopy merged images of OT1.0 (green channel) and the RFP channel (red, mCherry expression for OT-hM3Dq and tdTomato for no-hM3Dq). Right: responses of the OT1.0 sensor measured in ACSF (baseline), 60 nM DCZ, and 100 nM OT. Scale bars, 100 μm. **c**, **d**. Example OT1.0 traces (**c**) and peak change (**d**) in OT1.0 fluorescence; where indicated, DCZ or OT were applied to the slices. n=5 slices from 2 mice for OT-hM3Dq and n=4 slices from 1 mouse for no-hM3Dq. \*\*\**p*<0.001 and n.s., not significant.

**Extended Data Fig. 4:**
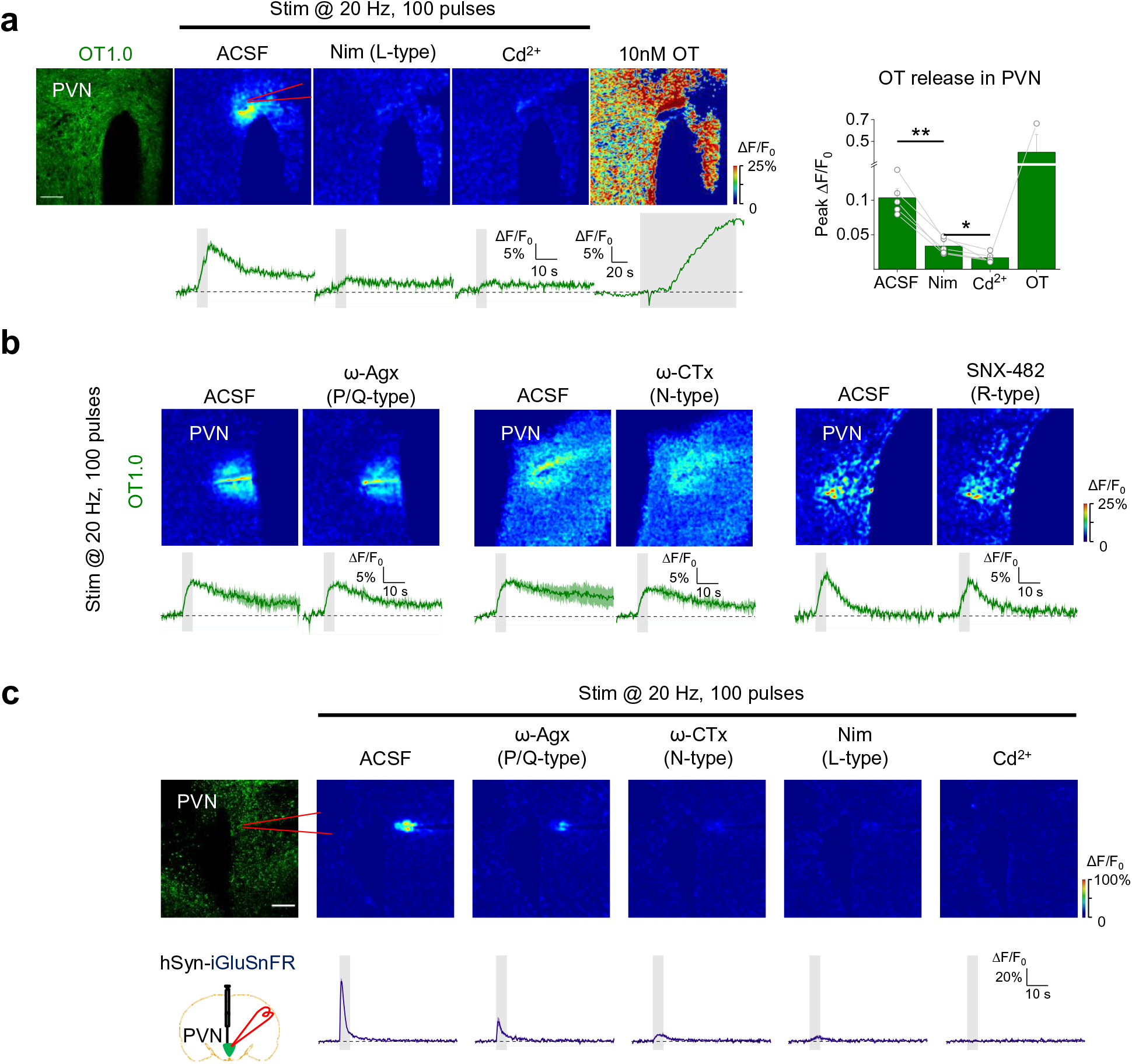
Dissecting the Ca^2+^ sources underlying somatodendritic OT release. **a.** Top left: representative image of OT1.0 expressed in the PVN (left). Also shown are example pseudocolor images (top row), corresponding traces (bottom row), and summary of the peak OT1.0 response (right) to 100 electrical stimuli delivered at 20 Hz in ACSF, nimodipine (Nim; 10 μM), Cd^2+^ (200 μM), or 100 nM OT. Scale bar, 100 μm. **b.** Representative pseudocolor images (top row) and corresponding traces (bottom row) of OT1.0 expressed in the PVN in response to 100 electrical stimuli delivered at 20 Hz in ACSF, ω-Agx-IVA (0.3 μM), ω-CTx (1 μM), or SNX-482 (100 nM) to block P/Q-, N-, and R-type VGCCs, respectively. Scale bars, 100 μm. **c.** Representative fluorescence image of iGluSnFR (top left) and schematic drawing depicting the experimental strategy (bottom left), related to Fig. 4e, f. Example pseudocolor images (top) and traces (bottom) of the change in iGluSnFR fluorescence in response to 100 electrical pulses delivered at 20 Hz in ACSF, ω-Agx-IVA (0.3 μM), ω-CTx (1 μM), nimodipine (Nim; 10 μM), or Cd^2+^ (200 μM) to block P/Q-, N-, L-type or all VGCCs, respectively (in this example, the same slice was sequentially perfused with the indicated blockers). Scale bar, 100 μm. \**p*<0.05 and \*\**p*<0.01.

**Extended Data Fig. 5:**
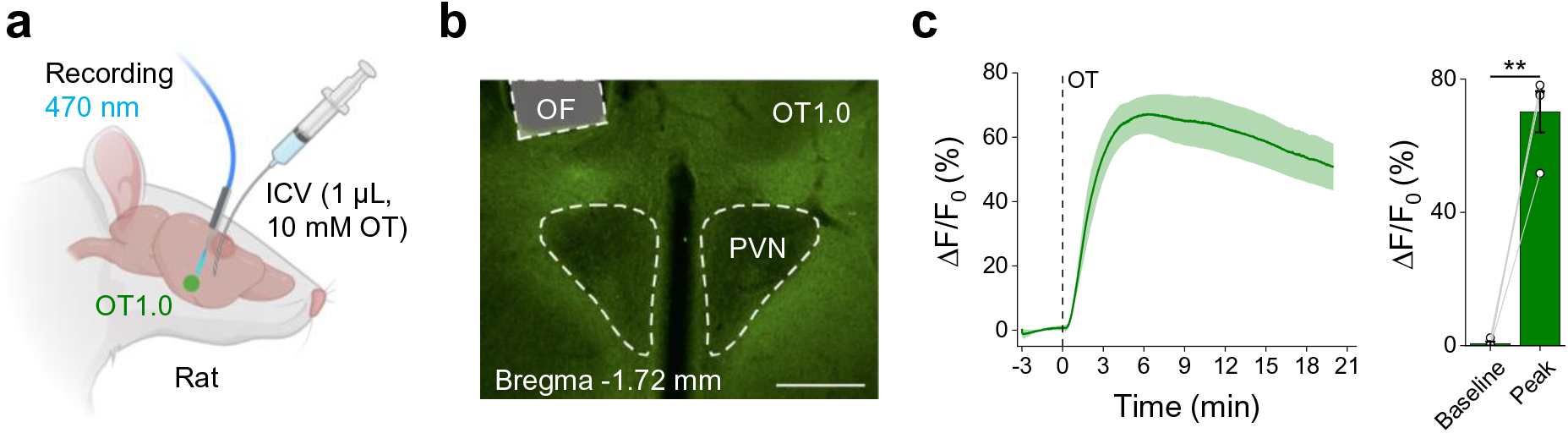
OT1.0 can detect intraventricularly injected OT in the PVN of rats. **a.** Schematic diagram depicting the experimental strategy for in vivo recording of OT1.0 in rats. An AAV expressing hSyn-OT1.0 was injected into the PVN of WT Sprague Dawley female rats; optical fibers were placed in the above PVN 2 weeks later, 10 mM OT (1 µL) was injected into the lateral ventricle during recording, and 470-nm light was used to excite the OT1.0 sensor together with isosbestic control signal (405 nm). **b.** Exemplary histological verification of the optic fiber placement and the OT1.0 expression in the periPVN area. OT1.0 was stained with anti-GFP antibody for visualization. OF, optic fiber. **c.** Average trace and quantification of OT1.0 signal. The OT1.0 and isosbestic signals were sampled at 1 Hz. n=4 rats. \*\**p*<0.01.

**Extended Data Fig. 6:**
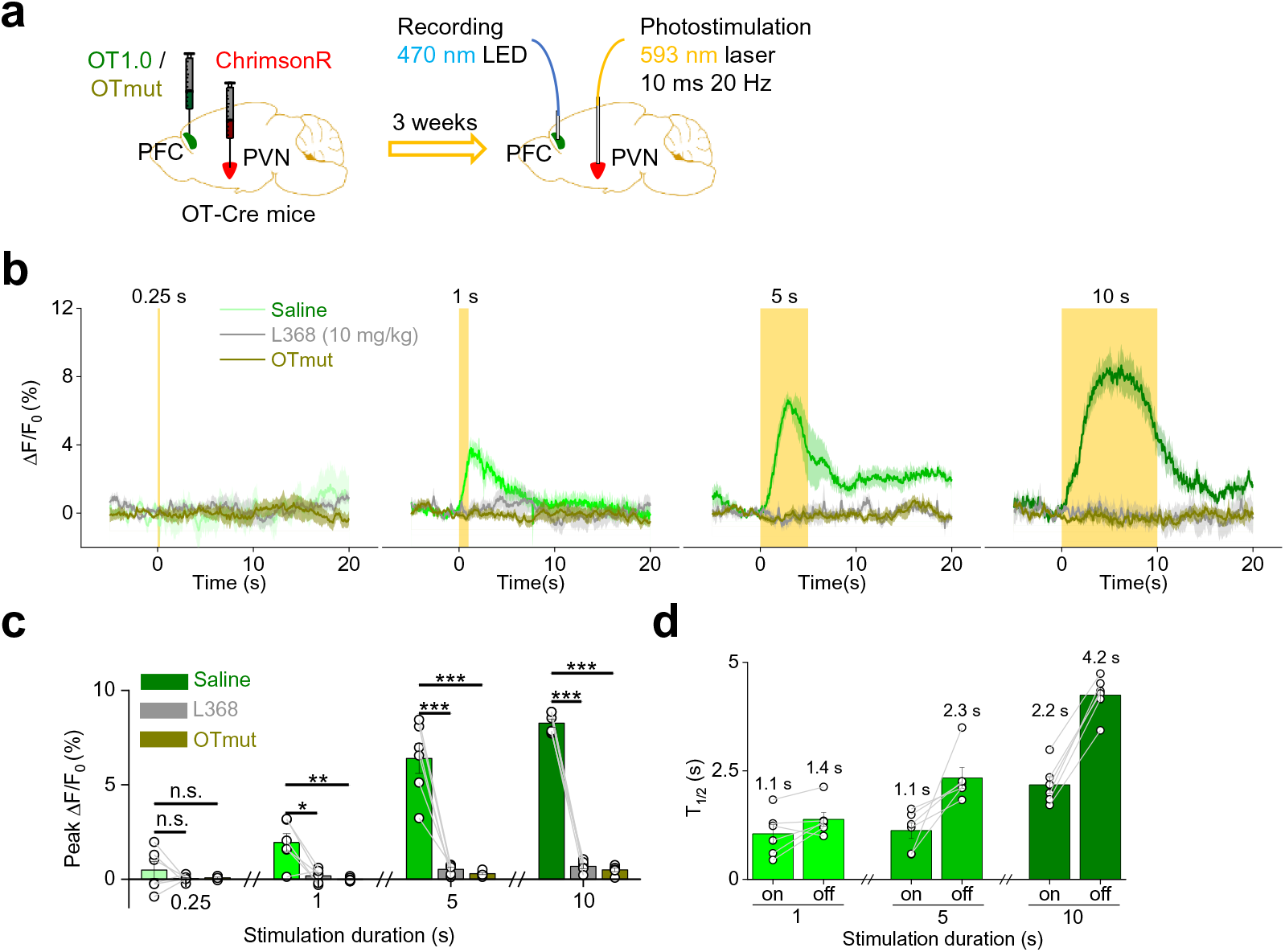
Optogenetic activation of neurons induces axonal OT release *in vivo* in freely moving mice. **a**. Schematic illustrations depicting the optogenetic activation experiments. An AAV expressing CAG-DIO-ChrimsonR-tdTomato was injected into the PVN, and an AAV expressing hSyn-OT1.0 (green) or hSyn-OTmut (gray) was injected into the mPFC of OT-Cre mice; 3 week after injection, optical fibers were implanted in the mPFC and PVN. **b**, **c**. Representative traces (**b**) and summary (**c**) of the peak change in OT1.0 (green) and OTmut (gray) fluorescence in mice that received an i.p. injection of saline or L368. Where indicated, a 593-nm laser delivered 10-ms pulses at 20 Hz for a duration of 0.25 s, 1 s, 5 s, or 10 s. Traces show the average of 5 individual trials in the same mouse. In **c**, n=6 mice per group. **d**. Summary of the rise time and decay time constants (T_1/2_) of the OT1.0 response to optogenetic activation with 10-ms pulses at 20 Hz for 1 s, 5 s, or 10 s; n=6 mice per group. \**p*<0.05, \*\**p*<0.01, \*\*\**p*<0.001, and n.s., not significant.

**Extended Data Fig. 7:**
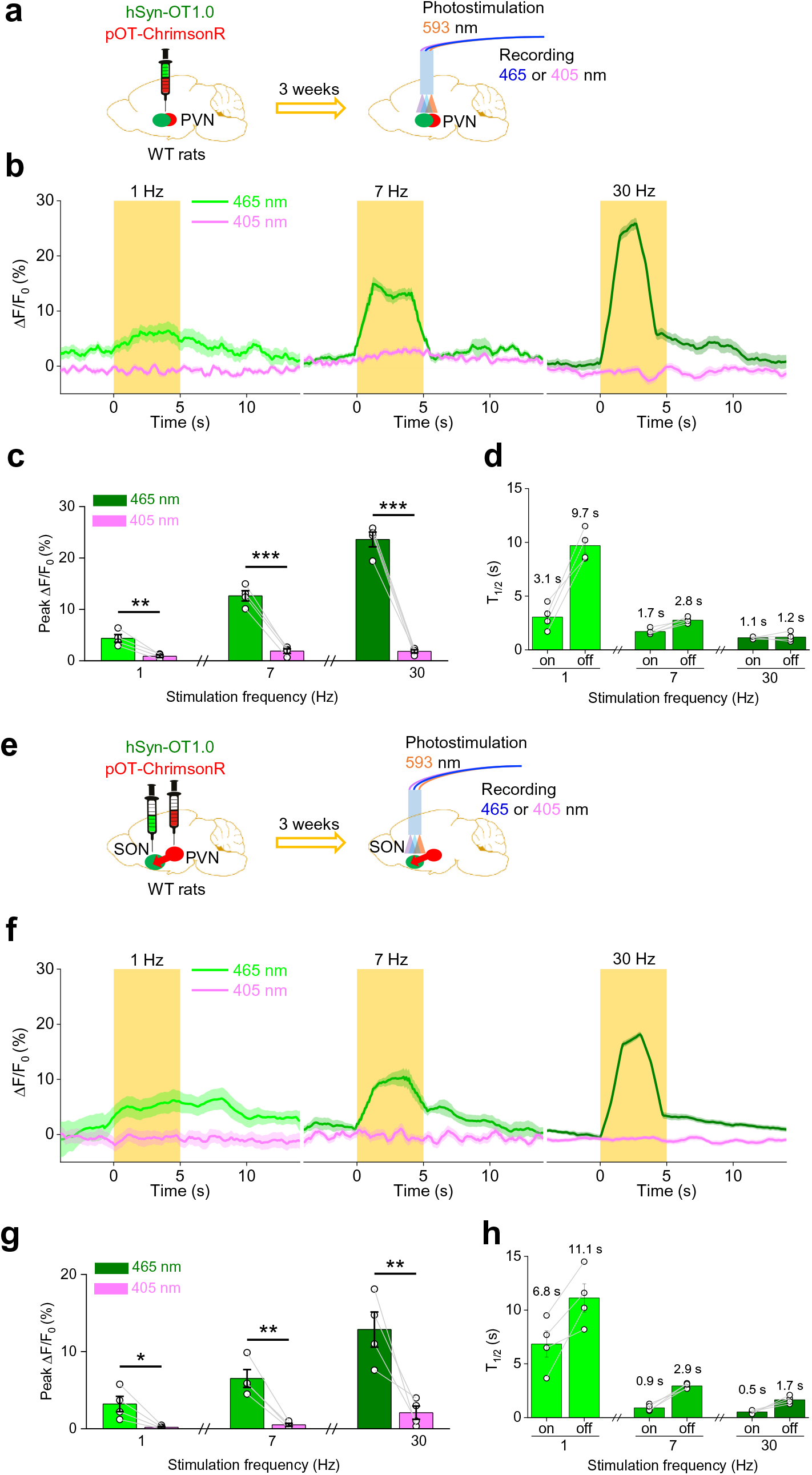
Optogenetic activation of neurons induces somatodendritic and axonal OT release *in vivo* in freely moving rats. **a.** Schematic illustrations depicting the optogenetic activation experiments with both the OT sensor and the optogenetic stimulation by ChrimsonR in the PVN. **b.** Representative traces recorded in the rat PVN of changes in normalized fluorescent emission ΔF/F_0_ (525 nm) during excitation at isosbestic control (405 nm, in purple) or sensor wavelength (465 nm, in green) before, during and after stimulation of ChrimsonR with pulses (at 593 nm, in orange) of 10 ms at a frequency of 1, 7 or 30 Hz for a total duration of 5 s). **c.** Summary of the peak changes in OT1.0 fluorescence emission (at 525 nm) during excitation at the sensor wavelength (465 nm, in green) or isosbestic control (405 nm, in purple) in rats PVN (at 1, 7, or 30 Hz photostimulation). **d.** Summary of the rise time (“on”) and decay time (“off”) constants (T_1/2_) of the OT1.0 response to photostimulation. In (**c**, **d**), n=4 rats per group. **e.** Schematic illustrations depicting the optogenetic activation experiments with OT sensor expressed in the SON and the optogenetic stimulator ChrimsonR expressed in the PVN. **f.** Representative traces recorded in the rat SON of changes in normalized fluorescent emission ΔF/F_0_ (525 nm) during excitation at isosbestic control (405 nm, in purple) or sensor wavelength (465 nm, in green) before, during and after stimulation of ChrimsonR (at 593 nm, in orange) with pulses of 10 ms at a frequency of 1, 7, or 30 Hz for a total duration of 5 s). **g.** Summary of the peak change in OT1.0 fluorescence emission (at 525 nm) during excitation at the sensor wavelength (465 nm, in green) or isosbestic control (405 nm, in purple) in rats SON (at 1, 7, or 30 Hz photostimulation). **h.** Summary of the rise time (“on”) and decay time (“off”) constants (T_1/2_) of the OT1.0 response to photostimulation. In (**g**, **h**), n=4 rats per group. \**p*<0.05, \*\**p*<0.01 and \*\*\**p*<0.001.

